# Background-matching patterns are attractive: support for a processing bias

**DOI:** 10.1101/2023.09.27.559753

**Authors:** Yseult Héjja-Brichard, Michel Raymond, Innes C. Cuthill, Tamra C. Mendelson, Julien P. Renoult

## Abstract

Background-matching visual patterns are selected to minimize detection by predators. However, once detected, what effects do these patterns have on conspecific perception? In humans, considerable evidence supports a "processing bias" by which people prefer images that match the spatial statistics of natural scenes, likely because the brain has evolved, or develops, to process such scenes efficiently. A direct but untested prediction of this bias is that people should prefer background-matching patterns compared to similar patterns that mismatch the spatial statistics of their surroundings. We conducted an online experiment to test this prediction on over 1700 participants. Our results show that a preference for the most frequent value of Fourier slope (a measure of spatial structure) observed in natural scenes can be modulated by an additional preference for background-matching. Because the initial stages of perception that underlie the efficient processing of visual scenes are widely conserved among animals, especially vertebrates, we expect this preference for background-matching patterns to be shared by other organisms. Two overlooked potential implications are that some animals may choose display sites with spatial statistics matching their own phenotype, and that the evolution of visual signals will, variously, be constrained or facilitated by such preferences for background-matching patterns.

**Highlights:**

- *Processing bias* aims to explain the origins of preference for sexual signals
- It predicts visual properties of background-matching patterns make them attractive
- An online experiment with over 1700 human participants confirms that prediction
- Results have important implications for the preference and design of animal signals

## Introduction

Understanding the origin of preferences for sexual signals is an ongoing focus of evolutionary biology. One hypothesis is that preferences originate as perceptual biases that arose in contexts other than mating (e.g., Endler & Basolo, 1998; Ryan & Cummings, 2013). “Processing bias” has been proposed as a pre-existing perceptual bias that could explain the origin of preferences and the subsequent evolution of sexual signals (Renoult & Mendelson, 2019). The processing bias hypothesis suggests that preferences originate as a bias for efficiently processed information (Renoult & Mendelson, 2019). Efficiency refers to the processing of information with both few errors and low metabolic cost, and it translates as the ease with which a signal is processed by the brain. Importantly, efficiency is known to increase the attractiveness of a stimulus, that is how likely that stimulus will be preferred when compared to another one (Fantz, 1957; Reber et al., 2004; Munar et al., 2015; Spehar et al., 2015; Renoult et al., 2016), probably because efficient information processing generates a feeling of fluency that is experienced as pleasant and therefore rewarding (Geller et al., 2022).

For visual signals, efficiency can be achieved by matching signal design with the sensitivity (i.e., tuning) of visual neurons (Barlow, 1961). A well-supported finding in neuroscience is that the visual system is adapted to efficiently process spatial statistics (e.g. luminance contrast distribution, feature orientations) of the environment (Field, 1987; Olshausen & Field, 1997; Simoncelli & Olshausen, 2001). As a result, stimuli that resemble the underlying spatial patterns of natural scenes will be processed more efficiently and experienced as pleasant or attractive. Indeed, a preference in humans for the statistical features of natural scenes is a robust finding in experimental psychology (e.g., Hagerhall et al., 2004; Spehar et al., 2003, 2015). In nature, background-matching is a type of camouflage whereby the colour and pattern of an animal’s phenotype mimic those of its surroundings (see e.g. Cuthill, 2019; Merilaita & Stevens, 2011). Given that background-matching relies on matching visual statistics of a stimulus with those of the habitat, the processing bias hypothesis predicts that background-matching patterns are visually attractive. This study aims to test this hypothesis and to discuss potential consequences of this prediction for the evolution of attractive and aversive signals in humans and other animals.

The attractiveness of patterns that mimic natural spatial statistics may seem to contradict predictions and empirical findings in visual ecology, which show that the function of this type of design is to go unnoticed. However, these two phenomena are not contradictory if we consider the detection of a stimulus and the processing of the information it contains as two distinct and sequential perceptual stages (Chun & Potter, 1995). Indeed, the same feature can function to minimize perceptual efficiency for detection, for example by reducing conspicuousness, but also to maximize processing efficiency once that pattern is detected. For instance, although the leopard’s rosettes were likely selected to increase their camouflage (Allen et al., 2010), those same rosettes are salient cues used by macaques to recognize the leopard as a predator (Coss et al., 2005). From an ecological perspective, this dichotomy explains why and how animals can bypass the trade-off between sexual and natural selection that is typically inferred in the context of detection, where sexual selection favours conspicuousness and natural selection favours crypsis. Animals can be both cryptic and conspicuous, for example if detection is only possible via private communication channels among conspecifics (Cummings et al., 2003), the appearance of the pattern is distance-dependent (Barnett et al., 2017), or if the signaller’s behaviour involves ostentatious displays, as in the Indian peafowl (*Pavo cristatus*), the timing of which can be controlled to avoid detection by predators (Kane et al., 2019). The question is then: what is the effect of a background-matching pattern on the attractiveness of an object or stimulus, once that stimulus has been detected?

There are two potential mechanisms by which the attractiveness of a stimulus can be influenced by its match with natural spatial statistics. A first hypothesis predicts a global preference for spatial statistics that are characteristic of natural visual scenes. That prediction was supported by Spehar and colleagues (2015), who found a strong relationship between human visual sensitivity and preference for simple visual patterns that mimic the spatial properties of natural scenes. Moreover, a large body of results in empirical aesthetics show that visual artwork often imitates natural spatial statistics (Graham & Field, 2008; Graham & Redies, 2010), increasing their perceived attractiveness. A second hypothesis suggests preferences will be stronger for background-matching patterns. For instance, Menzel and collaborators (2015) varied the slope of the background against which faces were presented and asked participants to rate their attractiveness. Faces were rated as more attractive when they were presented against a background that was closer to the statistics of both natural scenes and faces, i.e. when faces had statistical properties in common with backgrounds.

Here, using psychophysical experiments with humans as a model, we provide empirical evidence that the level of background-matching is correlated with the level of visual attractiveness, once a stimulus is detected. We used abstract stimuli made from white noise images to ensure that measured responses were due to the processing of low-level visual statistics as performed by the perceptual system, and not to cognitive mechanisms involving object recognition. This approach reduces the effect on preference of any benefit or danger associated with the stimulus, thereby facilitating the interpretation of results within the framework of processing bias. In total, we have designed three tasks. In the first task, we used a detection task to assess the effectiveness in minimizing detectability of different circular patterned stimuli (‘targets’) that varied in their level of background-matching. The backgrounds were artificial but with spatial statistics either similar to those of many natural scenes or manipulated away from typical values. Participants were required to click on the target as quickly as possible. This first task was necessary to demonstrate that our method for manipulating stimuli results in perceived variation in background-matching. In the second and third tasks, we evaluated the attractiveness of these targets, using the same patterned stimuli as for the detection task but removing the detectability constraint. Participants were asked in a forced-choice task to indicate which of the two patterns they preferred. In the second task, stimuli were presented in pairs against a uniform grey background to assess the baseline attractiveness of the target stimuli, independent of the background. This task measures the absolute preference for our stimuli, that is, without any background-matching effect. In the third task, stimuli were presented in pairs against the same patterned backgrounds as for the detection task, so that the degree of background-matching varied across stimuli, but stimuli were outlined to make them easily detectable. The third task, therefore, tested our main prediction that background-matching patterns are attractive when rendered detectable.

## Methods

### Online experiment and participant recruitment

The experiment took place online on a website hosted by the Montpellier Bioinformatics Biodiversity platform (University of Montpellier, France) and is available for demonstration purposes at https://isemsurvey.mbb.univ-montp2.fr/pattern/ The experiment was programmed using a custom HTML script (including JavaScript, CSS, and PHP) and jsPsych, a JavaScript library for creating online experiments (de Leeuw, 2015), and is available to play on all standard internet browsers. The experiment had to be conducted with a desktop or laptop computer, excluding smartphones and tablets.

### Stimulus design

Sets of stimuli for the three tasks were generated with the Python programming language (Python Software Foundation, https://www.python.org/). The procedure to create each image for the different experimental conditions for the first task was the following: two randomly generated white noise images (650 x 650 pixels), a target image and a background image, were first created and then Fourier transformed to change the slope of the relationship between the log-power spectrum and the log-frequency (i.e., the “Fourier slope”) corresponding to our different conditions (background image: slope = -1, -2, -3; target image: slope = background slope ± 0.25; ± 0.5; ±0.75). Concretely, an image contains features that span different scales, from broad (e.g. the homogeneous blue coloration of the sky) to fine details (e.g. sharp contrasts caused by grass). Those features have different spatial frequencies, low frequencies for broad-scale features and high frequencies for fine details, which are processed separately by the visual system (Musel et al., 2013). Measuring the Fourier slope of an image allows us to quantify the relative distribution of those spatial frequencies, which is known to influence visual preferences (see e.g., Nguyen & Spehar, 2021).

To manipulate the Fourier slope of our white noise images, we first converted our image to the Fourier domain using the Fast Fourier Transform function implemented in Python. After normalizing its Fourier transform by its greatest absolute value, we separated the image into a phase matrix and an amplitude matrix. We then estimated the rotational average of the power spectrum for each frequency before estimating the slope of the power spectrum with a linear regression. To create an image with the desired slope, we first generated a residual matrix from the difference between the image and an even amplitude spectrum with the desired slope. We then applied the residuals to an amplitude matrix with the desired slope and applied it to the phase matrix before reconstructing the image with an inverse Fourier transform. The obtained output image now has the desired slope value. To estimate the Fourier slope of an image, we fast Fourier transformed it and computed the rotational average of the power spectrum for all spatial frequencies between 10 and 110 cycles per image. This frequency selection limits edge effects and high-frequency noise. To give equal weight to all frequencies, we also binned (n=20) each power spectrum before estimating the slope value of the averaged power spectrum using a linear regression.

A negative linear relationship on a log-log plot indicates that the structure of a pattern tends to be scale-invariant and most natural scenes are characterized by a Fourier slope close to a value of -2 (Pouli et al., 2013). However, methods of Fourier slope calculation vary across the literature, and thus any comparison of slope values across datasets must be based on the same calculation. We therefore used our method of Fourier slope calculation to estimate the slope of 4,319 images of the Landscape Pictures dataset (Rougetet, 2020), which contains images of different types of landscapes (e.g. mountain, beach, desert), to confirm the value of the dominant slope in natural scenes. We found an average slope of -2.39 (SD: 0.31) for those images.

To generate stimuli, a circular selection with a radius of 80 pixels was selected from a random location of a target image and superimposed onto a background image at the same random coordinates. Targets could appear anywhere on the background, as long as they did not get cropped out of the frame (see Figures 1 and 2A for several examples). To eliminate pixelated edges, a very slight amount of blur was applied to the target edges. More specifically, a Gaussian filter (with a standard deviation of 0.5) was applied around the edge of each target before superimposing it onto the background image. Similarly, Gaussian filtering (with the same standard deviation of 0.5) was applied to the background image at the target’s location to smooth the contours around the target. This procedure had only a marginal effect on the slope of the power spectrum of the entire targets/backgrounds. The final shape and dimensions of those stimuli were a square of 650 x 650 pixels. Those dimensions are fixed and do not vary with the monitor size of participants.

**Figure 1:**
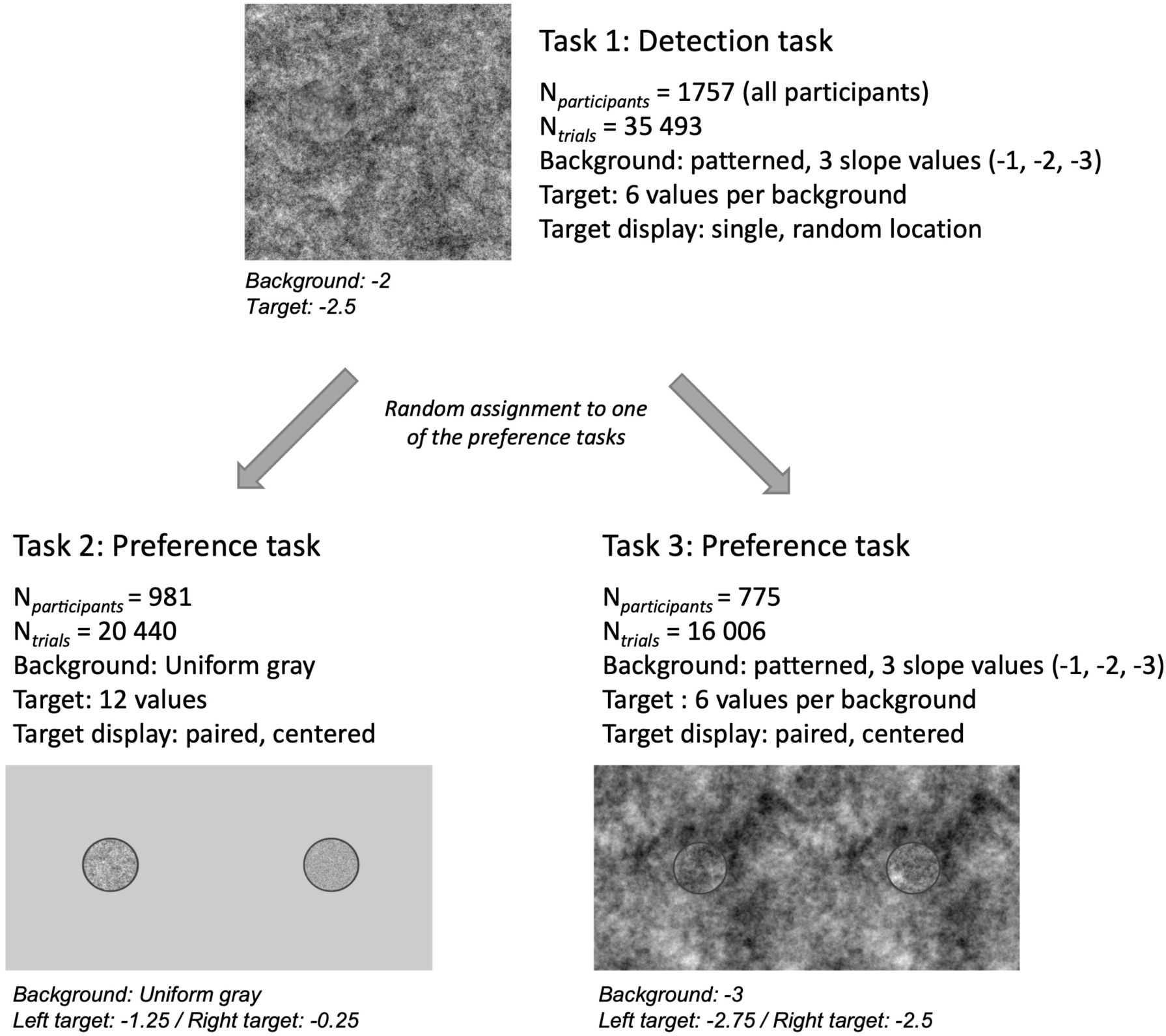
Overview of the experiment. For each of the three tasks, the number of participants that completed them and the total number of trials this represents are reported along with the main characteristics of the stimuli. An example of those stimuli is also included.

**Figure 2:**
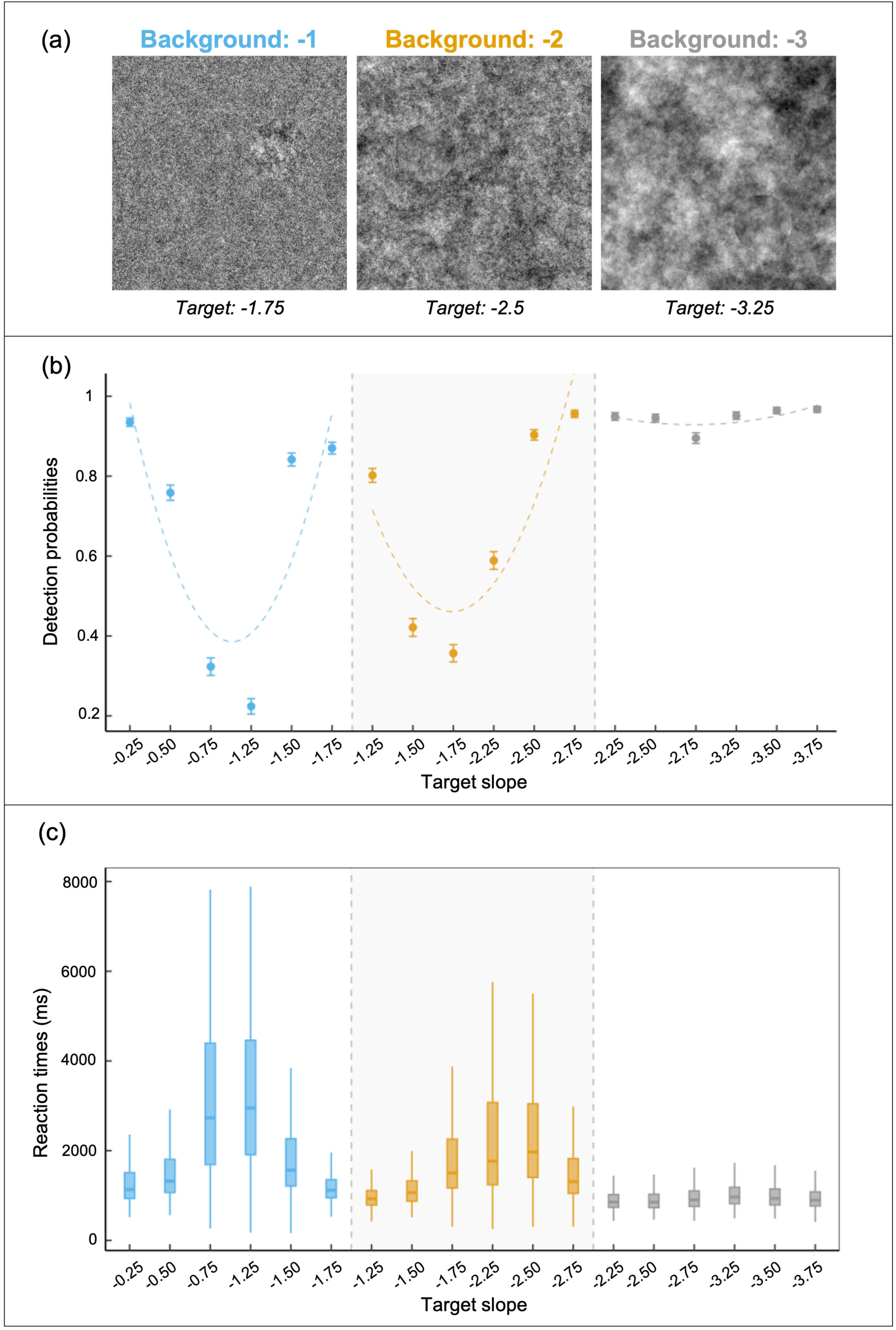
Detection task. (a) Examples of the presented stimuli with target values indicated below each image. The Fourier slope, always negative, is steeper for images with more low spatial frequency features, such as luminance variation extending over several pixels. (b) Detection probabilities for the different conditions, computed as the number of trials for which the target was detected divided by the total number of presented trials. Error bars correspond to 95% confidence intervals around the mean (represented by a full dot). Background slope values are color-coded: blue for -1, orange for -2, and grey for -3. Dashed lines represent a quadratic fit to the data for each background separately (-1, -2, and -3). Arrows indicate the corresponding background value. (c) Reaction times (ms) for detected trials of the different conditions. Box plots indicate the amount of within-condition variation in reaction times. Background slope values are color-coded: blue for -1, orange for -2, and grey for -3.

Using the same procedure, for the second and third tasks (forced-choice tasks), we created a second set of stimuli of larger dimensions (650 x 1300 pixels) to contain two targets, one centred in each half of the stimulus (Figure 3A and 4A). To make the targets conspicuous (i.e. easily detectable), we outlined them with a dark grey contour (linewidth: 1.5 and pixel intensity of the grey colour: 64). The set of stimuli for the second task was created with a uniform, grey background (pixel intensity of the grey colour: 128) in contrast to the patterned background of the third task. Here, the colour properties of the grey contour (i.e. target outline) are given on a uint8 scale where a pixel intensity value of 255 corresponds to white and 0 to black.

**Figure 3:**
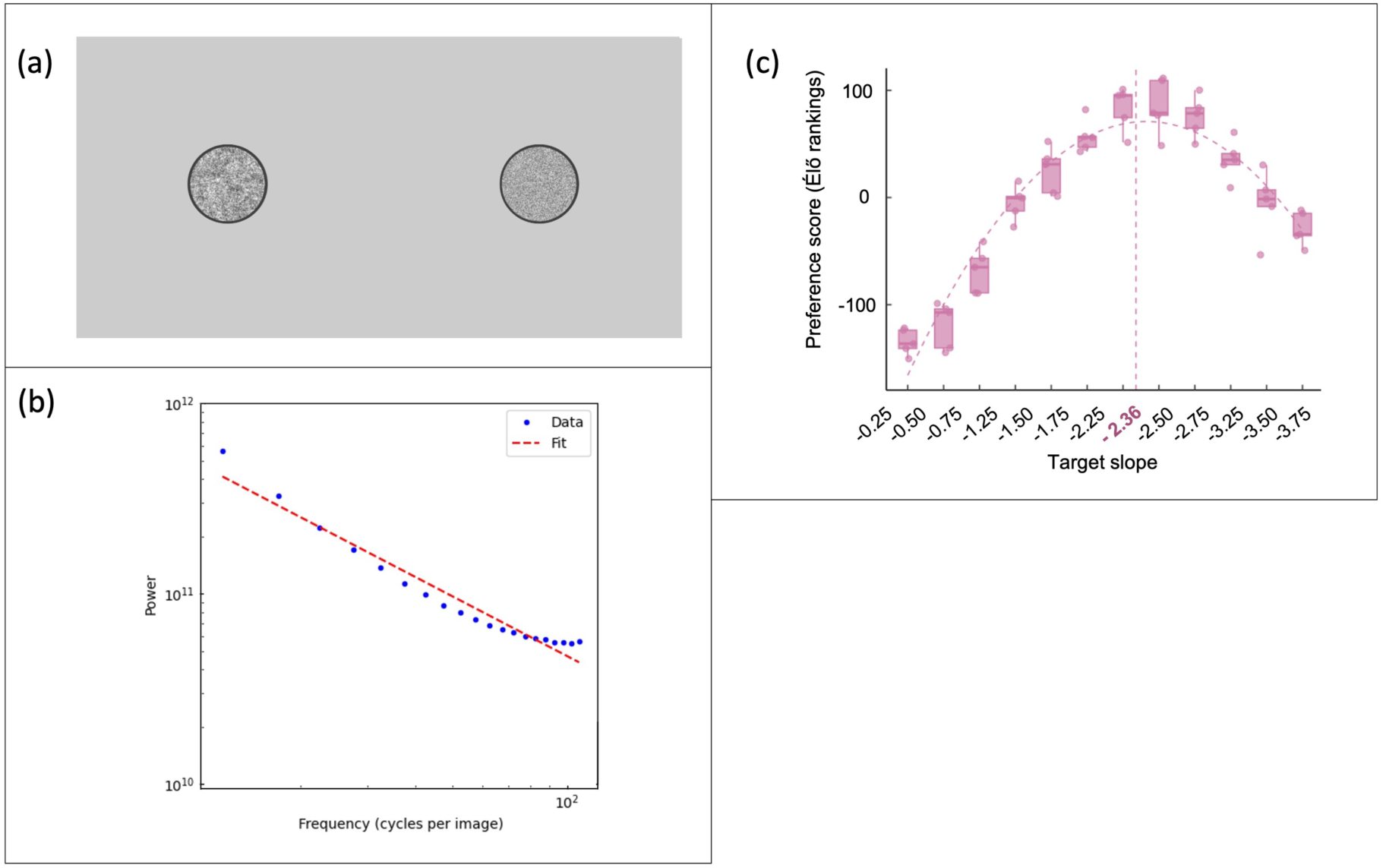
Preference task on grey background. (a) Example of a presented stimulus. Target slopes are -1.25 for the left target and -0.25 for the right target. In this example, we expected participants to prefer the left target. For display reasons, the background color here is lighter than that of the stimuli used in the experiment (see Methods). (b) Measurement of the typical Fourier slope of natural images. Log-log plot of the averaged power spectrum from which the Fourier slope of an image is derived (red regression line), here representing the averaged Fourier slope of images from the Landscape Pictures dataset. (c) Élő rankings for the two-alternative forced choice (AFC) task with a grey background. Each condition (i.e. target slope value) is represented by five closed circles (made more visible using jittering) corresponding to five different stimuli versions. Dots indicate the mean ranking values for each of the five stimuli versions of each condition (indicated on the x-axis). Box plots give the variability of preference scores across those versions. The boxplot central line indicates the median, the hinges the 25th and 75th percentiles, and the upper (lower) whiskers extend from the hinge to the largest (smallest) value no further than 1.5 * IQR (inter-quartile range, or distance between the first and third quartiles) from the hinge. The dotted line indicates a quadratic fit of the data.

**Figure 4.**
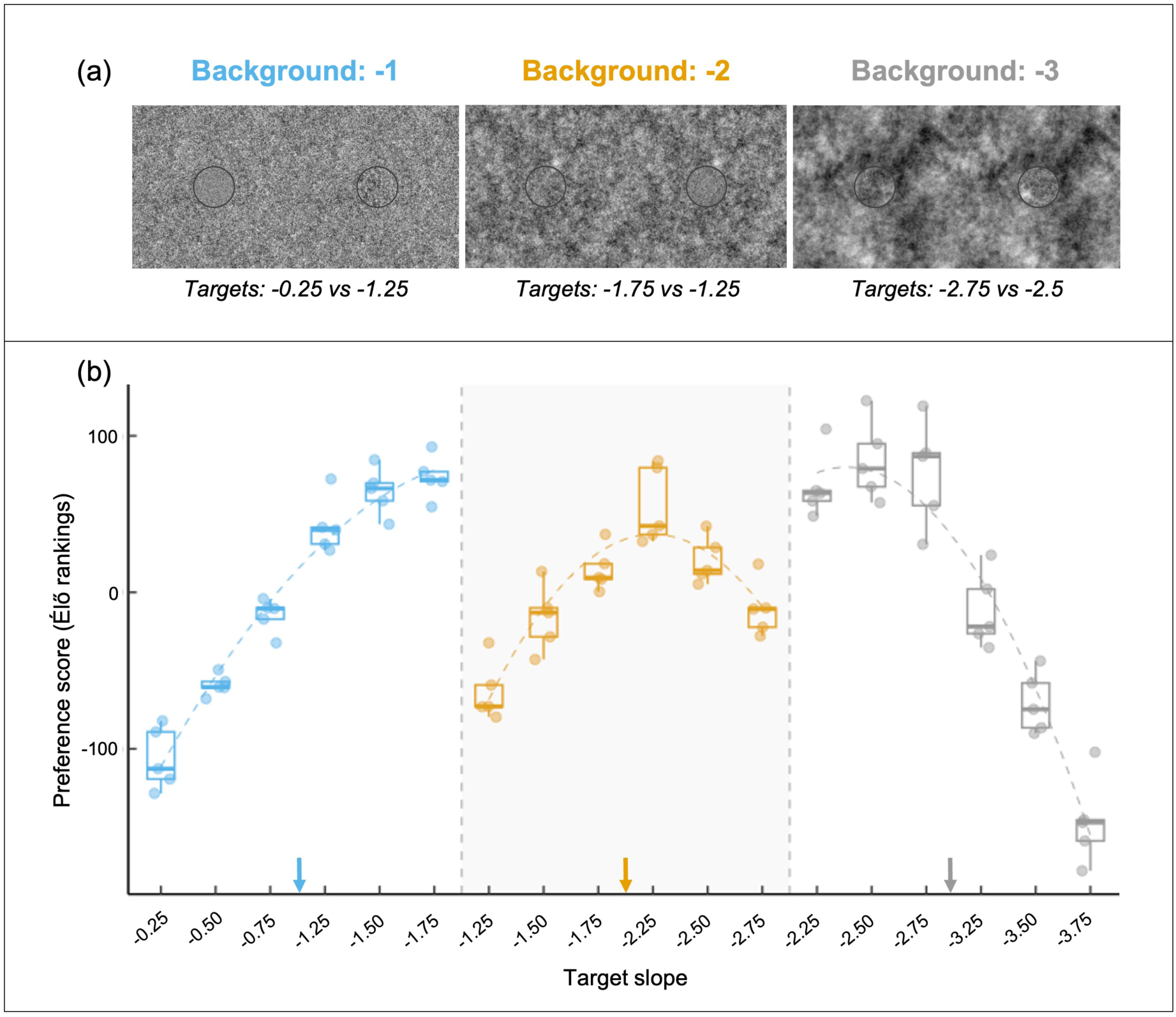
Preference task on a patterned background. (a) Examples of the presented stimuli with target values indicated below each image. (b) Élő rankings for the 2-AFC task with a patterned background. Each condition (i.e. one combination of background slope and target slope) is represented by five closed circles (made more visible using jittering) corresponding to five different stimulus versions. Box plots give the variability of preference scores across the 5 stimulus versions. The boxplot central line indicates the median, the hinges the 25th and 75th percentiles and the upper (lower) whiskers extend from the hinge to the largest (smallest) value no further than 1.5 * IQR (inter-quartile range, or distance between the first and third quartiles) from the hinge. Background slope values are colour-coded: blue for -1, orange for -2, and grey for -3. Dashed lines represent a quadratic fit to the data for each background separately (-1, -2, and -3), and arrows indicate the corresponding background value.

For each of the three tasks, we created 5 versions of stimulus images per condition to introduce variability in the pattern design. The slope value of all five versions was identical, but the resulting patterns varied in their local pixel arrangement. Thus, in total, task 1 had 18 different conditions (3 levels for the background variable: -1; -2; -3 and 6 levels for the difference between the background and the target slope: ± 0.25; ± 0.5; ± 0.75), each with 5 versions of the stimulus images, representing a total of 90 unique images. Task 2 had a total of 78 conditions (1 background slope value: “grey”, 2 targets with 6 slope differences each: ±0.25; ±0.5; ±0.75, and 2 sides: left or right), each with 5 stimulus versions, resulting in 390 unique images. Task 3 with a patterned background had a total of 90 conditions (3 background slope values: -1; -2; -3, 2 targets with 6 slope differences each: ±0.25; ±0.5; ±0.75, and 2 sides: left or right), each with 5 stimulus versions, resulting in 450 unique images. Note that paired targets were never identical.

### Experimental design

Each participant engaged in two tasks: the detection task (“Task 1”) and one of the two forced-choice tasks (grey or patterned background; “Task 2” or “Task 3”), which was randomly assigned. The presentation order of the two tasks was counterbalanced between participants (Figure 1).

### Detection task (Task 1)

Each stimulus was displayed at the centre of the webpage, one at a time, in a randomized order for a maximum duration of 8000 ms (see Figure 2A for stimulus examples). The intertrial interval was picked from a uniform distribution between 500 and 2000 ms (in steps of 1 ms) to avoid habituation (see e.g. Geer, 1966). The task contained 21 trials, which correspond to the number of images seen by each participant. This number was kept low to ensure that participants would complete the experiment. Each stimulus was randomly picked from the pool of 90 created images and presented only once, except for three of the first ten trials that were repeated at the very end to check for within-participant consistency. At the end of each trial (i.e. after the participant clicked on the stimulus with a mouse or a laptop’s touchpad, or when the response time limit had been reached), participants received feedback in the shape of a yellow circle, highlighting the location of the target. Before the test trials, three practice trials were presented to make sure the participant understood the task. Participants were instructed to “Click on the target as fast as possible”. The location and time of mouse clicks were recorded and used to define detection probability and response times. Detection probability was computed as the number of trials for which a target was correctly detected (i.e., clicked on) divided by the total number of trials. Response times correspond to the recorded time of a mouse click and could take any value between 0 and 8000 ms (maximum presentation time). In the absence of a mouse click, the response time of a trial was set to NA.

### Forced-choice task (Task 2 or 3)

Stimuli were presented in a randomized order for a maximum duration of 12 000 ms. The intertrial interval was picked from a uniform distribution between 500 and 2000 ms (in steps of 1 ms). The task contained 21 trials, which correspond to the number of images seen by each participant. Each pair of stimuli randomly picked from the pool of 390 (grey background, Task 2) or 450 (patterned background, Task 3) created images was presented only once, except for three of the first ten trials that were repeated at the very end to check for within-participant consistency. Pairs of stimuli differed in two ways: the background slope values (-1; -2; -3 for task 3 or a uniform grey background for task 2) and the position (left-side or right-side of the image) of the target could take four and two different values, respectively. Importantly, both targets always differed in their slope value. Participants were asked the following question, displayed above the stimuli: “Compare the patterns in the two circles, which one do you prefer?”. The location of participants’ mouse clicks was recorded and used to determine the preferred pattern.

Preference scores for a target with a specific slope value were expressed as a ranking, computed via the Élő score (using the EloChoice package in R, v0.29.4; Neumann, 2019), an iterative algorithm traditionally used in chess games and that performs pairwise comparisons. Élő scores are now frequently used as a measure of aesthetic value and are well suited for unbalanced designs as is the case in our study (see Tribot et al., 2018; Clark et al., 2018 for similar use; Goodspeed, 2017 for a comparison of different pairwise ranking systems). Briefly, each pair of stimuli is similar to a contest where the preferred stimulus wins points, and the non-preferred stimulus loses the same number of points. Importantly, Élő rankings are a relative ranking system, meaning that stimuli with higher Élő scores have a higher probability of being preferred (i.e. of being perceived as attractive) than stimuli with lower Élő scores. Moreover, Élő scores can only be compared across contests that have a common stimulus configuration. In our experiment, this means that the background against which targets are presented must be identical for targets that are being compared. Thus, for task 3, Élő scores were computed separately for each background (i.e., -1, -2, and -3), because pairs of targets were presented against the same background, and thus inter-background comparisons are not possible. As in Clark and collaborators (2018), we used the elochoice function with initial Élő scores set to 0, and *k* set to 100. We ran 1,000 simulations to make sure ratings were stable. We ran the analysis image-wise, that is, by considering each of the five versions of a stimulus as an independent condition to account for potential variability between stimulus versions. Running the same code but by considering all five versions of a stimulus as the same condition did not yield different results. To ensure we had sufficient trials to obtain stable Élő scores, we estimated the minimum number of participants required to achieve stability in ratings, that is, how many participants need to have rated the stimuli, for the estimated ranking of those stimuli to no longer change. This is an important consideration given the incremental nature of the ranking algorithm. To compute a reliability index, we used the raterprog function of the EloChoice package with the ratershuffle argument set to 10 to avoid a participant order effect. We found that using the data from 378 participants for images with a grey background (reliability index = 0.57, Figure S1) and from 263 participants for images with a patterned background (reliability index = 0.59, Figure S2) was sufficient to reach stable ratings (reliability index of 0.6 is interpreted as indicating “good reliability”, see Neumann, 2019).

### Questionnaires

Participants were asked 5 questions before and 2 questions after they performed both tasks. Questions were demographic in nature and related to education level and frequency of exposure to visual art (Table S1).

### Statistical analyses

All statistical analyses were performed using the software R (R Core Team, 2021, version 4.4.1). We removed participants who did not finish the experiment and participants who took part in the experiment more than once (IP address verification procedure, as described in Supplementary Text S1). In total, we kept the data of 1757 participants out of the 2567 participation events that were recorded (68.6% of participants). For details on sociodemographic data, see the supplementary material (Supplementary Text S2).

For the detection task, we removed 49 trials (0.14% of trials) where responses were faster than neurophysiological limits (i.e. below 150 ms, see e.g. Thorpe et al., 1996) and therefore likely correspond to accidental clicking. For the two-alternative forced-choice tasks, we found no lateral bias in participants’ responses: the left target was chosen in 50.5 to 51% of the trials.

In each task, three of the 18 trials (i.e. corresponding to different stimuli) were presented twice, resulting in a total of 21 trials. Responses to the second presentation of those three repeated trials were compared to those from the first presentation by performing paired comparisons with t-tests. This procedure allows for checking within-participant consistency without dramatically lengthening the experiment. For the detection task, we found no difference for the detection probability, as defined by the number of times the stimulus target was correctly located divided by the total number of presentations of the stimulus (*t* = - 0.8238, df = 177.16, *P* = 0.4112). We found the response time for the second repetition (mean: 1,577) to be faster than for the first repetition (mean: 1,733) (*t* = -6.7704, df = 13147, *P* < 0.0001), suggesting participants were improving in this aspect of the task. For the preference tasks, we found no difference between the first and second presentation of the three repeated trials in terms of preferred stimulus, neither for the grey background (*t* = 0.5223, df = 7754, *P* = 0.6015), nor for the patterned background (*t* = 0.2833, df = 6034, *P* = 0.7770). Further analyses for all tasks were conducted on the non-repeated trials only.

For the detection task, we first assessed detection success by fitting participants’ responses with a linear mixed-effects binomial model. The response variable is binary, with 1 corresponding to a detected target and 0 to a missed target. To investigate variation in response times across conditions, we fitted a second generalised linear model with a gamma distribution for the target-detected trials only. For both models, independent variables included the slope of the background, the absolute difference between the target slope and the background slope, and the interaction of both variables. Three random effects were also added: the stimulus displayed (*1|Stimulus*), the condition tested (*1|Condition*), and the participants’ anonymised identifier (*1|Participant*).

For the preference tasks, we fitted the data with linear mixed-effects models (*lme4* package, Bates et al., 2015) using maximum likelihood estimation. We first estimated the Élő score for each image and used those scores as the dependent variable. For Task 2 (with a grey background), independent explanatory variables (fixed effects) included the slope value of the target and the quadratic of this variable, shown to be needed when comparing models’ outputs. We added the stimulus version (5 versions per stimulus type) as a random effect. There were in total 60 different images (that is, across conditions and for each stimulus version separately). For Task 3 (with patterned backgrounds), we ran three similar models for each background separately. Independent explanatory variables (fixed effects) included the absolute difference between the background slope and the target slope, the sign of the difference (positive or negative, coded as 1 and -1, respectively), and their interaction. Similarly to Task 2, we ran additional models with the slope value of the target and the quadratic of this variable to estimate the peak of the preference. Again, we added the stimulus version (5 versions per stimulus type) as a random effect. There were in total 30 different images per background (that is, across conditions and for each stimulus version separately).

In addition to the Élő score approach described for Tasks 2 and 3, we replicated the analyses using alternative methods based on linear mixed-effects binomial models. Data for those binomial models correspond to the individual participants’ responses. For both tasks, these models yielded qualitatively similar results (see Supplementary Text S3, Tables S2, S3, and Figure S3).

### Ethical note

Participants were recruited through advertisements on dedicated mailing lists and on social networks. Participants were not paid, and they were required to be at least 16 years old. Approvals from the French (deemed exempt, CNIL number 2-21044) and American (deemed exempt, UMBC IRB #522) ethics committees were obtained before the start of the experiment (see Supplementary Text S1 for additional details). We therefore confirm that our work adhered to the ethics of scientific publication as detailed in the Ethical Principles of Psychologists and Code of Conduct (https://www.apa.org/ethics/code/index). We also certify that the research was conducted in accordance with the ethical principles of the 1964 Declaration of Helsinki and its later amendments.

Before launching the online experiment, the study’s objectives and planned analyses were pre-registered following open science practices (available on https://osf.io/m7ap6/).

## Results

### Task 1: Detection task

The goal of the first task was to validate that our method effectively generated perceptually meaningful variation in background-matching. We predicted that targets with a Fourier slope closer to the slope of the background would be less frequently detected and require more time to detect. Results confirmed this prediction: targets with slopes more similar to the background (smaller slope difference value) were the hardest to find (lower detection probabilities and higher response times, Figure 2B and C, Tables S4 and S5, Figures S4 and S5). Targets presented against the steepest background slope (BKG -3) were easier to detect compared to other background conditions (BKG -1 and BKG -2) (Figure 2B). This is likely because on backgrounds with a Fourier slope of -3, more of the spectral power (contrast) is concentrated in lower spatial frequencies: the pattern of dark and light is more coarse-grained. As a result, phase-mismatches at the target’s edge are clearer than on finer-grained backgrounds (e.g. slopes of -2 or -1). In simple terms, on coarse-grained backgrounds there are larger areas of dark and light so, even if the target has the same coarse-grained pattern, there is a greater chance of light areas on the target lying on, or next to, dark areas of the background, and likewise for dark on light. Nevertheless, our results confirm that, at least with BKG -1 and BKG -2, we successfully generated perceptually meaningful variation in background-matching across stimuli.

### Task 2: Forced-choice task to assess preference for natural statistics in abstract visual patterns

Our second task tested whether the patterns we generated for the targets in the first task also varied in their attractiveness against a grey background. Targets with differing Fourier slopes elicited markedly different preferences (Figure 3C). The resulting Élő score rankings can be fitted with a simple quadratic model (ranking ∼ slope + slope²), which predicts a preference peak for a target slope of -2.36, with 95% CIs [-2.33, -2.39] (fit adjusted R²: 0.9136, Table S6). This preference peak is similar to the preferred peak value reported in previous studies (see e.g. Spehar et al., 2015) and also to the dominant Fourier slope of the natural visual environment, which is typically between -1.81 and -2.4, depending on the study (Pouli et al., 2013). As noted above, when computing the Fourier slope of a large dataset of landscape pictures, we found an average Fourier slope of -2.39 (SD: 0.31), a value very similar to the preference peak inferred from Task 2 (-2.36). Thus, our results further corroborate the general finding that people prefer images with the Fourier slope of natural scenes. Combined with those from the detection task, our results validate the use of the Fourier slope to manipulate both the effective level of background-matching and the attractiveness of stimuli. They also indicate a baseline pattern preference that must be accounted for when considering a preference for background-matching stimuli in Task 3.

### Task 3: Forced-choice task to assess the attractiveness of background-matching patterns

In our third task, we assessed the attractiveness of patterns that varied in their degree of background-matching. Because they were clearly outlined, all targets were assumed to be equally conspicuous, and we assessed participants’ preferences for different levels of background-matching with a two-alternative forced choice (2-AFC) task.

Running separate linear mixed-models for each background to investigate a background-matching effect, we found similar trends for backgrounds -1 and -3. We found a significant interaction between the absolute slope difference value (background - target) and the sign of that difference (see Tables S7, S8, S9 for statistical details), meaning that preference increases with greater background-matching (smaller difference between the background slope and the target slope) when the sign of the difference is negative, but decreases when the difference is positive (Figure 4B).

For background -2, the absolute slope difference value and the sign of that difference are both independently significant (no interaction). Here, preference increased with greater background-matching, and preferences were higher when the difference between the slope of the target and the background was positive.

Those results, especially for backgrounds -1 and -3, suggest the presence of an effect in addition to background-matching.

We thus tested for a preference for natural statistics in Task 3, which would translate as a preference for targets with a slope closer to -2.39. We ran separate quadratic models for each background (ranking ∼ slope + slope²) to estimate the target slope value with the highest ranking against each background (see Tables S10, S11, S12). For background -1, preference peaked at a target slope of -2, with 95% CIs [-1.71, -2.7]. For background -2, the preferred target slope was -2.14, with 95% CIs [-2.11, -2.18]. For background -3, it was -2.48, with 95% CIs [-2.61, -2.24]. Thus, confidence intervals for backgrounds -1 and -3 included the average Fourier slope value of natural scenes (-2.39), indicating that this well-established preference only appears when presented against certain backgrounds.

Confidence intervals for background -2 did not include the average Fourier slope of natural scenes. Indeed, there was no overlap between the confidence intervals for backgrounds -2 and -3, indicating different preference peaks for these two backgrounds. The fact that preferences peaked at different maxima depending on the background, and that these peaks were shifted away from -2.39 toward the value of the background, confirms that background-matching is an important factor in shaping pattern preferences.

## Discussion

Our study aimed to test a prediction of the processing bias hypothesis, which proposes that animals prefer efficiently processed stimuli. Because brains are tuned to the statistical patterns of natural scenes (whether through evolution or life experience), stimuli that mimic those patterns should be efficiently processed and therefore, in some sense, “attractive” (Redies et al., 2007; Spehar et al., 2015). Background-matching is one way in which organisms mimic the spatial statistics of natural scenes; therefore, a processing bias hypothesis predicts that background-matching patterns should be attractive when detected. We developed an online experiment to test that prediction, by estimating preferences for stimuli that varied in the degree of background-matching while holding conspicuousness constant. Our results constitute two main findings. First, they corroborate previous studies showing a clear preference for the average Fourier slope of natural scenes (Task 2). Second, they demonstrate that this preference can be modified, shifting toward the pattern of the background (Task 3). Whether and how a preference for natural scene statistics drives the evolution of animal signalling is discussed.

### Evidence for a processing bias

Results of the first task demonstrated our ability to generate variation in background-matching across stimuli, by manipulating the Fourier slope of abstract patterns. Stimuli in which the Fourier slope of the circular target more closely matched the slope of the background were located less frequently by participants and took longer to detect. Our results thus align with previous studies showing a link between Fourier slope and background-matching effectiveness. For instance, performing a spectral analysis of octopus patterns, Josef and colleagues (2012) found that octopuses adapt their background-matching pattern to specific features of their environment by imitating the Fourier slope of nearby structures. In cuttlefish, the Fourier slope of an animal’s pattern better matches its background when hiding from predators or prey compared to when signalling (Zylinski et al., 2011). Our results complement these correlational studies, showing that the Fourier slope can be manipulated to generate variation in the detectability of background-matching stimuli. We note that, in addition to the neural coding of the observer, absolute properties of the background also affect how easy it is to avoid detection through background matching. For example, a phase mismatch between target and background (manifested as a clear edge to the target) is harder to detect on backgrounds where more of the spectral power is at high spatial frequencies. In Figure 2, the lowest detection probabilities and longest response times are for background-matching targets on backgrounds of slope -1 (fine-grained) as compared to a background of -2 and particularly compared to the coarse-grained background of -3. In other words, a background-matching target on a fine-grained background is hard to detect wherever it lies, but a background-matching target on a coarse-grained background may, by chance, lie on or next to a region it mismatches.

Results of the second task corroborated a preference for the Fourier slope that is typical of natural scenes, based on a forced-choice task for stimuli of different slopes against a grey background (see also results with binomial models: Supplementary Text S3 and Table S11). A preference for a Fourier slope typical of natural scenes has been reported in numerous studies in experimental psychology (e.g., Hagerhall et al., 2004; Spehar et al., 2003); here we corroborate that finding with a much larger sample size (Figures 1 and 3). In a particularly compelling study, Spehar and collaborators (2015) showed that the Fourier slope typical of most natural scenes is not only the most attractive, but also the slope to which people have the highest visual sensitivity (i.e. absolute detection, discrimination, and contrast sensitivities). Their result emphasizes the potential influence of efficient processing on visual preferences: patterns whose properties coincide with visual sensitivity are processed more efficiently by the visual system and are generally preferred over patterns that are more dissimilar to visual sensitivity peaks. A preference for natural scene statistics is also consistent with the processing bias hypothesis (Renoult & Mendelson, 2019), and it can explain a large body of results in empirical aesthetics, which show that visual artwork imitates natural spatial statistics (Graham & Field, 2008; Graham & Redies, 2010). For example, faces painted by portrait artists across time and cultures tend to have a Fourier slope value approximating that of natural landscapes even though real faces typically do not, suggesting that artists increase the attractiveness of their work by unconsciously mimicking the spatial statistics of natural scenes (Redies et al., 2007).

Results of the third task also confirm a preference for target slopes near -2, that is, for the Fourier slope typical of natural scenes. However, estimated preference maxima also revealed that this preference is additionally modulated by the background slope, indicating a preference for patterns that more closely match the pattern of the background. Our results therefore support the processing bias hypothesis and also highlight a potential interaction between absolute and relative preferences for stimuli that match the spatial statistics of visual scenes. An absolute preference for natural statistics would be determined by the general, “hard-coded” adaptation of neuronal sensitivities to the average spatial statistics of natural scenes, or by the long-term tuning of neuronal sensitivities to these statistics during early development (Sengpiel et al., 1999); whereas, a relative preference, corresponding to a preference for patterns that match the local background, would be determined by the short-term, plastic tuning of neuronal sensitivity. Both preferences have been demonstrated in the literature (e.g. Menzel et al., 2015; Spehar et al., 2015); here, we find evidence for both in a single experiment. We found an absolute preference for patterns with the average Fourier slope of natural scenes and furthermore that this preference can be modified by the visual statistics of the background against which a stimulus is displayed.

### Implications for animal preferences

The extent to which these results observed in humans have implications for the ecology of signalling and preferences in other animal species primarily depends on whether a preference for natural statistics exists in non-human animals. The adaptation of the brain to efficiently process natural statistics was first shown in primates and later found in rodents (mice, ferrets: Akerman et al., 2004; Qiu et al., 2021), probably reflecting a general strategy shared across most animals (Baden, 2021; Laughlin, 1981; Troscianko & Osorio, 2023). Indeed, there are known similarities between the early visual systems of humans and those of macaques (Fize et al., 2011), cats (Kang et al., 2009), mice (B. Williams et al., 2021) and birds (W. Clark et al., 2022), and there is strong evidence that tuning of the visual system to the statistics of natural habitats allows efficient information processing in vertebrates (Olshausen & Field, 1997; Simoncelli & Olshausen, 2001) and insects (Laughlin, 1981). Moreover, a substantial body of research on non-human animals suggests a bias for efficiently processed stimuli (reviewed in Renoult & Mendelson, 2019). For example, stimuli with continuous or symmetric surfaces/lines, fractal structures, temporal repetition, or those that are representative of a perceptual category (i.e., prototypical) are both processed more efficiently and preferred by animals compared to stimuli lacking these characteristics.

Recently, Hardenbicker and Tedore (2023) discovered that peacock jumping spiders prefer images with the Fourier slope of natural scenes over two other options. In their study, preference was defined as time spent oriented towards presented stimuli, with greater time indicating greater preference. Inspection time is routinely used to assess preferences in several taxa, including humans, with studies in infants relying on a similar protocol (e.g., Langlois et al., 1991). Taken together, these findings strongly suggest that a preference for natural statistics likely occurs in non-human animals as well as in the participants in our study.

### Implications for animal signalling

If preferences can originate as pre-existing perceptual biases for background-matching patterns, this implies that signal patterns will be shaped by the spatial statistics of the environment, i.e., patterns that mimic those statistics will be efficiently processed and therefore attractive (Renoult & Mendelson, 2019). We are not claiming that the attractiveness of a visual scene, camouflage pattern or artwork is necessarily underpinned by the same emotional or physiological state as sexual arousal. A scene or stimulus may be attractive without evoking sexual arousal. The key point is that the valence in each case is positive, eliciting attention and a desire for physical proximity. In the case of sexually arousing stimuli, the animal seeks to mate; in the case of efficiently processed information, the animal seeks to continue processing the stimulus. Efficiently processed signals might also increase attractiveness through “misattribution”, whereby the observer misattributes the pleasure emerging from efficient processing to the observed object instead (Renoult & Mendelson, 2019). If these motivations are additive and increase the probability of mate choice, a preference for background-matching patterns could influence the direction of sexual signal evolution towards including a background-matching component (Mendelson et al., 2025).

This prediction of the processing bias hypothesis is an extension of ”sensory exploitation” (Ryan et al., 1990) and “sensory drive” (Endler, 1992), which propose that the environment shapes both sexual signals and preferences. Most examples of sensory drive focus on a bias for detectability, with animal signals evolving to be more easily detected (Endler & Basolo, 1998), and for particular colours. The African cichlid fish *Pundamilia nyererei,* for example, lives in an environment with red-shifted light. Females exhibit an increased expression of red-sensitive photoreceptors that render red stimuli more detectable, and males exhibit more reddish coloration (Seehausen et al., 2008). Our results suggest that expanding sensory drive beyond the framework of signal detection, to account for the effect of processing bias once a signal is detected, can help us understand the evolution of signal patterns in addition to coloration.

A preference for natural and background statistics in animals also has implications for signalling strategy. Results of our study suggest that courters should respond behaviourally or physiologically so that the statistical features of their patterns mimic not only those of the average natural statistics encountered by the population, but also potentially of the local background against which they are displaying. And if their sexual signal changes, as in some fish that change sex during development, individuals might choose an alternative displaying site that better matches their pattern. Previous studies have shown that animals choose their display site in relation to their own sexual signals (Endler & Thery, 1996; Fleishman et al., 2022); however, those studies focus on coloration, showing that animals gain attractiveness by maximizing contrast with the background and the ambient light. Our results shift the focus to signal patterns, but predict that for this signal component, animals could benefit from mimicking spatial statistics of the visual environment. At the very least, if signalling favours departure of patterns from that of the background (e.g., for detectability), the resulting reduction in signal attractiveness will need to be overcome. Importantly, matching its pattern with the background scene does not prevent an animal from being conspicuous: multimodal signalling and distance-dependence readily allow for the combination of ostentatious features (e.g., colours or sounds) with cryptic features (Marshall, 2000; Tullberg et al., 2005; Barnett et al., 2017). In general, our study aligns with earlier suggestions (Endler, 1992), recently brought back into focus (Caves et al., 2023), to consider the importance of the visual background in animal signalling strategies.

Another strategy to leverage the attractiveness of background-matching patterns might be seasonal variation in detectability. Individuals that are camouflaged most of the year could be temporarily rendered more attractive by the addition of a detectable feature during the mating season. Male darter fish (genus *Etheostoma)*, for example, exhibit vivid and species-specific colours during the breeding season that appear to complement rather than conceal their underlying background-matching pattern (Kuehne & Barbour, 1983; Page, 1983, Figure 5B). Indeed, females in at least two species of darters are attracted similarly to conspecific colour and pattern when these signal components are decoupled (Williams & Mendelson, 2011). An attractive and conspicuous seasonal addition could thus enhance, and be enhanced by, the underlying attractiveness of the background-matching pattern.

**Figure 5.**
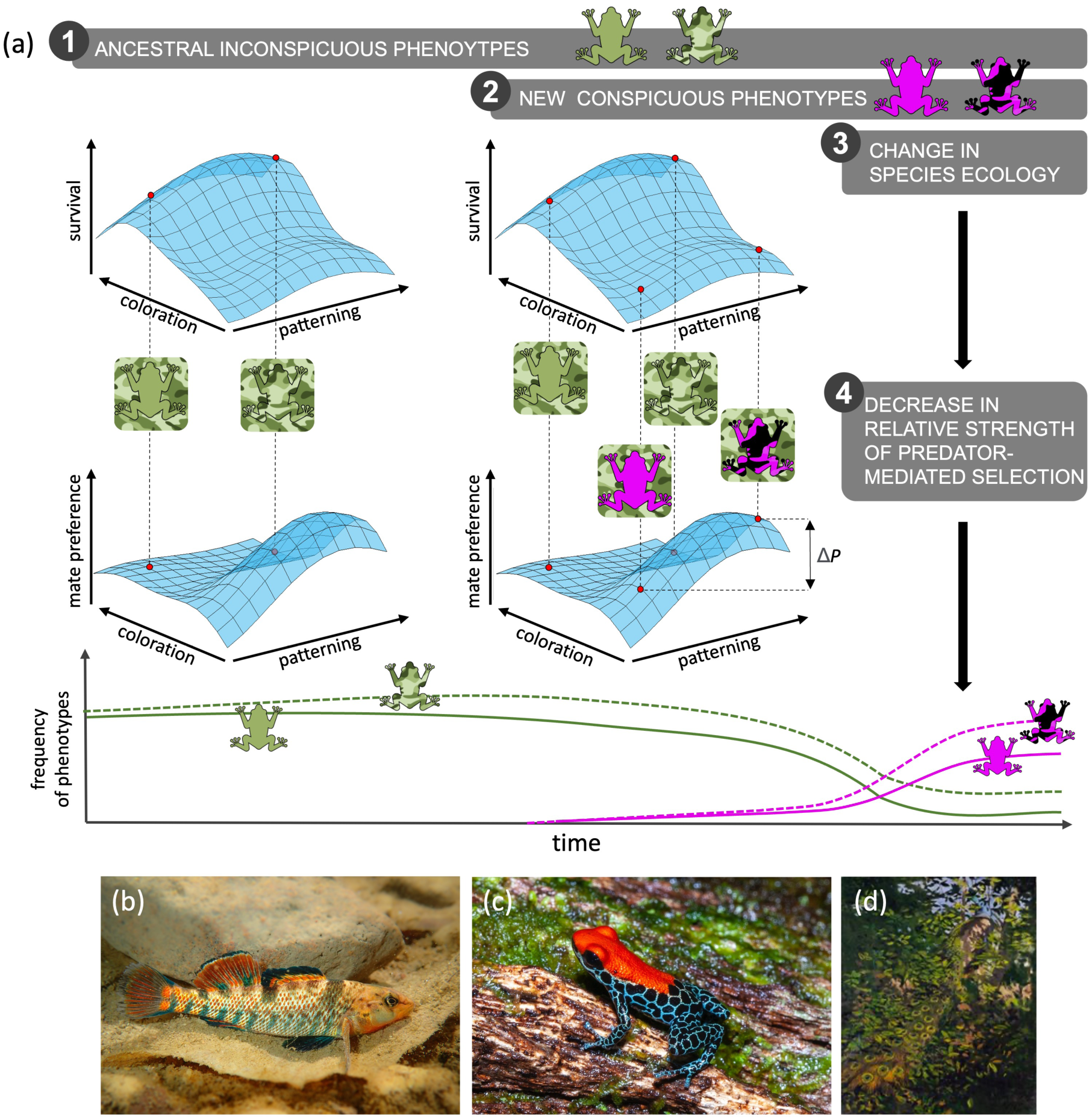
Evolution from background-matching to sexual signalling through the exploitation of processing bias. **Panel (a): 1.** Natural selection due to predation generally results in inconspicuous phenotypes. Phenotypes with a pattern that matches that of the environment (‘inconspicuous pattern’) are well-camouflaged and thus gradually increase in frequency in the population. Such phenotypes typically do not elicit strong preferences among conspecifics because of their low detectability. **2.** Any new mutation increasing conspicuousness should be negatively selected by predation (low survival). However, because they are highly detectable by potential mates and thus reduce their cost of mating, or if they enhance sexual arousal, these phenotypes are potentially attractive. Among conspicuous phenotypes, background-matching patterns that match the visual structure of the habitat (‘conspicuous patterned’) should be preferred (ΔP) over those that do not (‘conspicuous plain’) because they are even more efficiently processed. In this study, we have provided the first empirical evidence supporting this prediction. **3.** These ‘conspicuous patterns’ would remain negatively selected by natural selection until a change in ecological factors (e.g., colonization of habitats or regions with fewer predators, the appearance of new, alternative prey, predators’ focus on new sensory cues) makes sexual selection strong enough to counteract the effects of predation on phenotype evolution 4. Thus, conspicuous phenotypes that have evolved from background-matching patterns can be evolutionarily co-opted by sexual selection. **Lower panels:** Examples of background-matching patterns underlying conspicuous colour and patterns. (b) Etheostoma caeruleum (Julien Renoult). (c) Ranitomeya reticulata (DH Fischer). (d) Painting of a peacock (Pavo cristatus) hidden in a tree by Abbott H. Thayer. Thayer, a naturalist painter of the 18th century, explored the idea that the primary function of animal patterns is camouflage. Although widely criticized at the time, Thayer’s idea is consistent with the proposed framework, that even the most canonical nuptial patterns can evolve from, and continue to function at least in part as, camouflage. (Abbott Handerson Thayer, Richard S. Meryman, Peacock in the Woods, study for book Concealing Coloration in the Animal Kingdom, 1907, oil on canvas, Smithsonian American Art Museum).

Processing bias may also play a role in the evolution of defensive signalling. For example, aposematic signals are typically conspicuous and easily learned by predators, consistent with efficient processing (e.g., Green et al., 2018). Notably, many of these warning signals are also attractive to conspecifics of the opposite sex (Rojas et al., 2018), as in the strawberry poison dart frog *Oophaga pumilio* (Maan & Cummings, 2008) and the red postman butterfly *Heliconius erato* (Finkbeiner et al., 2014). Many aposematic patterns in frogs appear as irregular, rounded blotches, which could be due to developmental constraints, but which could also be perceived as colourful embellishments of attractive background-matching patterns (Figure 5C; Galindo Uribe et al., 2014). The attractiveness of aposematic patterns, therefore, may be due to the ease with which they are processed, with sexual selection and natural selection similarly aligned in the evolution of warning signals. However, an alternative way to dissuade receivers (e.g., predators), aside from the vivid coloration typical of aposematic signals, is to exhibit patterns that generate discomfort, such as patterns that depart from natural scene statistics. This is indeed what is found in the colour patterns of some Lepidoptera (Penacchio et al., 2024). Additionally, an aversion or ‘visual discomfort’ in response to patterns that depart from natural scene statistics has been found in humans for whom a simple deviation from the average Fourier slope predicts judgements of discomfort from images (Penacchio & Wilkins, 2015).

### The roles of natural and sexual selection in the evolution of courtship communication

The idea that background matching can be attractive may appear counterintuitive given the extensive literature documenting conflicts between natural and sexual selection in courtship communication. Because traditional approaches to understanding the evolution of signal design are based on signal detection, natural and sexual selection are usually pitted in conflict along an axis of detectability, with natural selection favouring crypsis and sexual selection favouring conspicuousness. Guppy fish highlight this trade-off: females prefer males with brighter and larger orange spots, but the size and colour of those spots depend on predator abundance (e.g., Breden & Stoner, 1987; Endler, 1978, 1980; Houde & Endler, 1990). However, while antagonism between these two types of selection is likely common when it comes to detection, this is not necessarily the case for all signal components, especially those processed once detection has occurred, such as the patterns in the current study. Our results suggest that natural and sexual selection can act synergistically for patterns: background-matching patterns may be favoured by natural selection because they reduce detectability to predators, and once detected by conspecifics, they also may contribute to enhance a signaller’s attractiveness and mating success. In this way, background-matching patterns could be evolutionarily co-opted by sexual selection. For example, if a change of ecological factors reduces predation pressure, a mutation that makes a signaller and its background-matching patterns more easily detectable, for example a conspicuous colour, could increase their attractiveness because of the efficiency with which the patterns are processed. Hidden preferences for those efficiently processed patterns thus are predicted to lead to an increase in the frequency of attractive phenotypes if ecological conditions permit (Figure 5).

An important limitation of these implications for natural and sexual selection is that predators and conspecifics often have different visual perception and viewing conditions. Fully characterizing the role of processing bias in the evolution of communication signals under different types of selection requires knowledge of the visual systems of interacting prey and predators. If they are sufficiently different, this reduces the conflict imposed by the two types of receivers, by facilitating the possibility of simultaneously cryptic (to predators) and conspicuous (to conspecifics) signals. On the other hand, these differences also mean that what is perceived as background matching by a predator may not be perceived as such by a potential mate. The distance at which predators and conspecifics view the signal should also be considered, given the known relationship between distance and conspicuousness or camouflage (e.g. Barnett et al., 2017; Marshall, 2000; Tullberg et al., 2005). The patterns examined in this study vary in their Fourier slope, and therefore in their degree of scale invariance, which makes the results relatively robust to small differences in viewing distance and visual acuity. Nevertheless, accounting for the specific psychophysical characteristics of the different receivers will be a crucial (and likely limiting) component for testing these predictions in realistic ecological contexts in animals. Although this might limit the generalization of our results, our study represents a proof of concept where we test the logic by which a bias for efficiently processed stimuli may influence the evolution of signalling strategies.

In sum, we provide robust evidence for a preference for patterns matching the average image statistics of natural scenes, and that this preference can be influenced by the pattern of the background against which a stimulus is viewed. Although demonstrated here only in humans, this result is expected to generalize to other species. By providing evidence of a processing bias, the key mechanism by which background-matching patterns can be co-opted to function as sexual signals, this study highlights the importance of considering the psychological landscape (Guilford & Dawkins, 1991) of signal receivers, and of interfacing research in evolutionary biology with the vast literature in human cognitive psychology.

## Data availability statement

A GitLab repository contains the code to reproduce the stimuli and the images used for the online experiment in high resolution, the pre-processed data and the code to reproduce the main analyses and figures: https://gitlab.com/yseulthb/camo_exp

## Acknowledgement

This work was supported by the National Science Foundation grant NSF IOS 2026334. We would like to thank Claudia Ximena Restrepo-Ortiz for her help with the Spanish translation of the online experiment. This work was supported by the National Science Foundation grant NSF IOS 2026334. We would like to thank Claudia Ximena Restrepo-Ortiz for her help with the Spanish translation of the online experiment. We are also grateful to Thomas Cronin, Mike Ryan, John A. Endler, and Mark Kirkpatrick for providing feedback on an earlier version of this manuscript. Finally, we thank the participants for taking part in our experiment.

## Supplementary material

### List of Supplementary Materials

**Figure S1:**
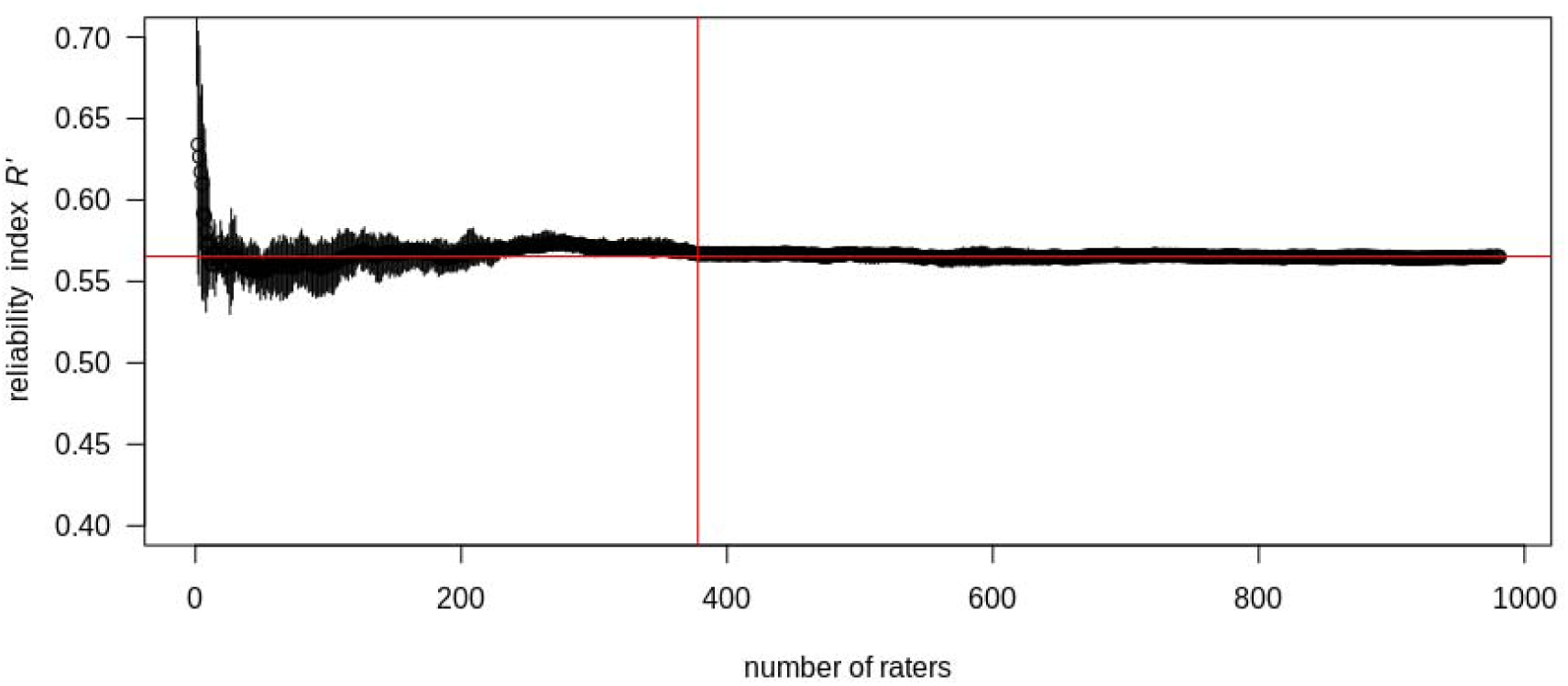
Ratings stability for the 2-AFC task with a grey background. The intersection of the two red lines indicates the number of raters needed to achieve the averaged reliability index (avgRI). AvgRI is measured as the average RI between 400 and 981 participants, for which we can see that ratings have achieved stability. The obtained avgRI is 0.567, corresponding to 378 raters and to a good reliability (Neumann and Clark, 2019). This was averaged across all 10 simulations, which correspond to a randomised order of raters.

**Figure S2:**
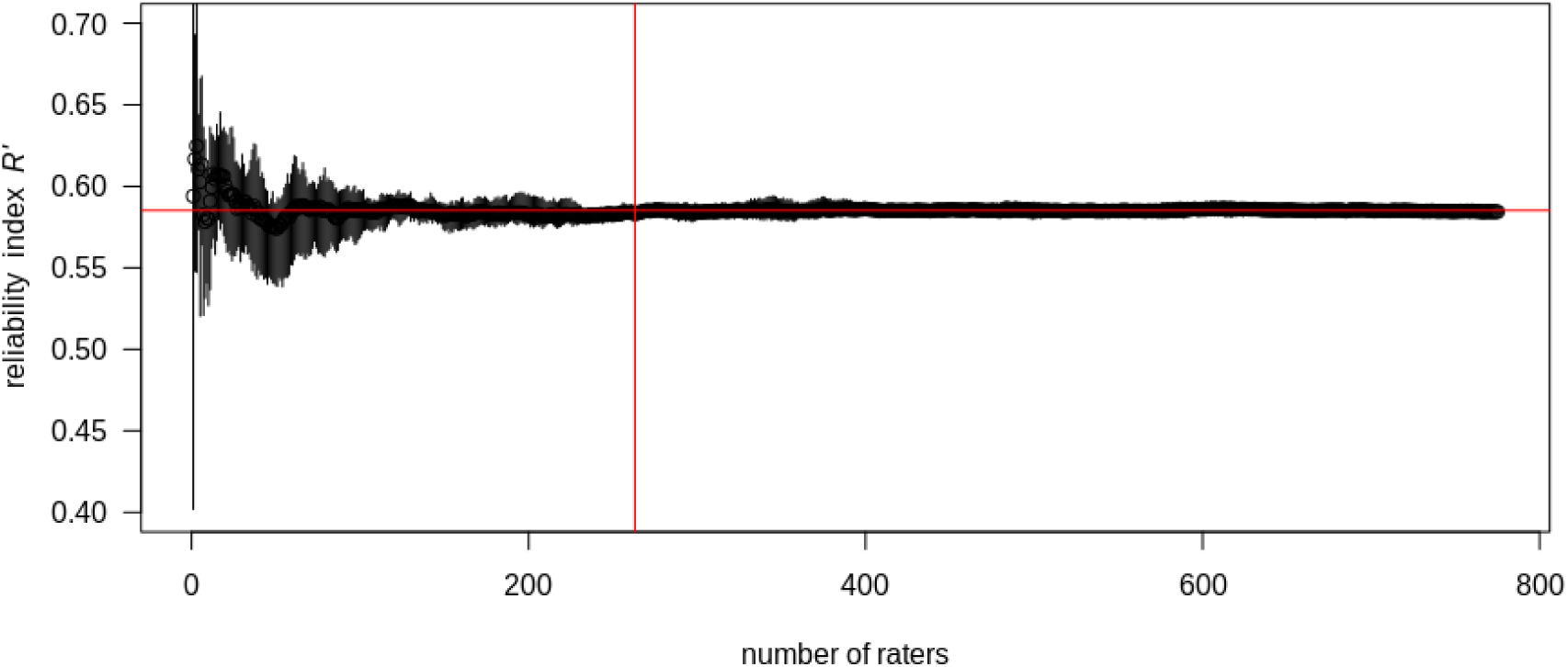
Ratings stability for the 2-AFC task with a patterned background. The intersection of the two red lines indicates the number of raters needed to achieve the averaged reliability index (avgRI). AvgRI is measured as the average RI between 250 and 775 participants, for which we can see ratings have achieved stability. The obtained avgRI is 0.585, corresponding to 263 raters and to a good reliability (Neumann and Clark, 2019). This was averaged across all 10 simulations, which correspond to a randomised order of raters.

**Table S1:**
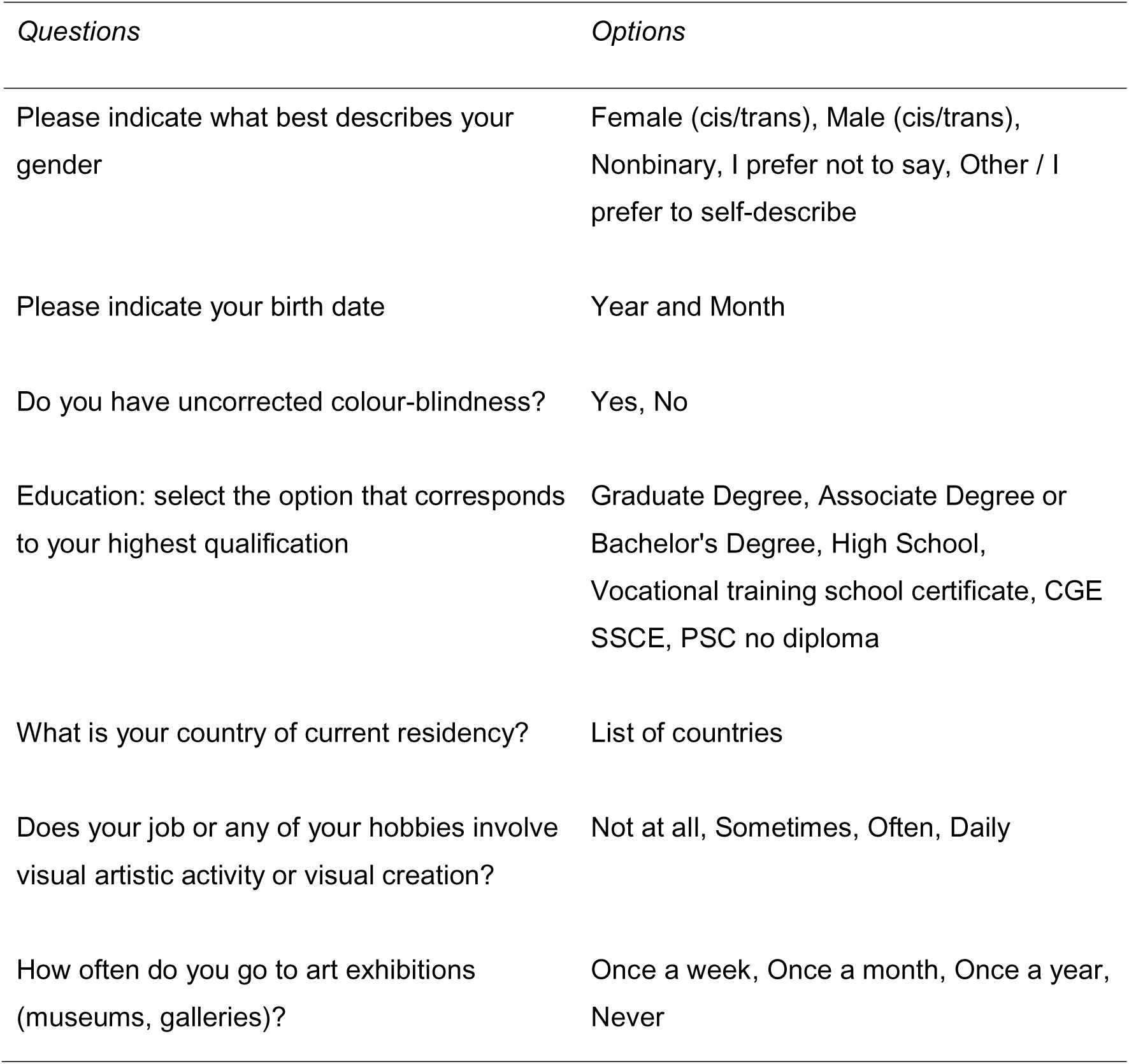
Experiment questionnaire and response options.

### Supplementary Text S1: Participants’ recruitment and procedure

**Recruitment**: Participants were recruited through advertisements on dedicated mailing lists and social networks. Participants were not paid, and they had to be at least 16 years old, corresponding to the legal digital majority in the EU. Before the experiment, participants were asked to report their month and year of birth to enforce that rule. For American citizens aged between 16 and 18 years, participants were asked to obtain their parents’ verbal approval before they participated in the experiment. Participants were asked to take part in the study only once. On top of informing them of that condition before the experiment. The IP address of the participant was only used to enforce that rule and was not associated with personal information, following a protocol approved by the CNIL. Collected data were stored on-site.

All research was performed following relevant guidelines/regulations. By French legislation, the protocols for this study have been submitted and approved by the French National Commission on Informatics and Liberty (CNIL number 2-21044). The CNIL, an independent public authority affiliated to the European Data Protection Board, is responsible for ensuring that information technology ethically serves the citizen and that it does not infringe on human identity, human rights, privacy or individual or public liberties. The present research is an online experiment with volunteers interacting remotely and anonymously, thus no ethical approval was required—nor possible—from an academic or scientific comity under French legislation, as their recourse is only possible if it is compulsory. The European GDPR (General Data Protection Regulation) was fully applied, and all participants were informed of the subject of the study.

**Procedure**: At the beginning of an experiment, participants were offered to choose the instruction language between French, English, and Spanish. After agreeing to the study details (consent form) and answering some questions about participants’ demographic details (Table S10), the experiment started.

### Supplementary Text S2: Sociodemographic data

Participants whose data were included in the analysis were 61% women (n=1069), 36% men (n=629), and 3% in other categories (non-binary: 38, prefer to self-describe: 5, prefer not to say: 16). A majority of participants had graduate degrees (66%, n=1162). 35 participants self-reported as colour-blind, representing less than 2% of the participants. Most participants were born after 1980, with a peak in the early 2000s, corresponding to undergraduate students who took part in the study as a way to get extra credit. When asked about their hobbies, most participants (82%) reported having hobbies that don’t involve visual creativity or only rarely. Most participants go to art exhibitions once a year or less (82%). Participants were mostly located in France (46%) and in the US (29%), with an additional good representation of Western Europe and Canada. Our participants are therefore WEIRD (from Western, Educated, Industrialized, Rich and Democratic countries as described in Henrich et al., 2010), which was expected given our advertising methods (mostly academic or professional mailing lists in Western countries).

It is possible that certain demographic traits, as well as higher exposure to visual art or creativity, might influence stimulus preferences. Therefore, we looked for potential relationships between demographic variables (age, gender, education level) and the ranking scores, and between visual exposure to art (presence of visual arts activity in jobs or hobbies, and frequency of art exhibition visits) and the ranking scores. For each demographic variable of interest, we computed the ranking score for patterned and grey images for a subset of the data, corresponding to the targeted participants. We then randomly sampled from the whole participant dataset a subset of the same size and computed the ranking for those participants as well (for both patterned and grey images). After making sure the rankings of both subsets (or groups of participants) were normally distributed, we compared them using a t-test. Overall, no difference was found for any of the tested sociodemographic variables.

### Supplementary Text S3: Statistical analyses with binomial models

In addition to the Élő score approach described in the manuscript for tasks 2 and 3, we replicated the analyses using alternative methods based on linear mixed-effects binomial models. Data for those binomial models correspond to the individual participants’ responses. A description of each of the included variables and of the model predictions is included for each task, followed by the tables of results.

**In task 2,** the goal was to test for a general preference for natural statistics. The measure of the average Fourier slope value of the Landscape dataset, which comprises 4319 pictures of various landscapes from different categories (e.g. beaches, forests), revealed that this average is at -2.39, consistent with the range of values usually reported for similar pictures (Pouli et al., 2013). This value was used as a reference when assessing a preference for natural statistics.

A linear mixed-effects binomial model was run on the data. The response variable *bino_click* was defined as a binary variable where 1 indicates that the left target was chosen (i.e. clicked on) and 0 that the right target was chosen. For each side (left and right), we calculated the Manhattan distance between the slope target and -2.39, to quantify “how similar to natural statistics the target is”. The explanatory variable was then calculated as the difference between Manhattan distances of the two sides: *[abs(slope_targetL-2.39) - abs(slope_targetR-2.39)]*. We predicted a preference for targets that have a Fourier slope closer to the one found in natural scenes (-2.39). This translates as a negative estimate for that explanatory variable: if the left target slope value is closer to -2.39 than the right target, the difference will be negative, while bino_click should tend towards 1 (click on target left). Three random effects were also included: the stimulus displayed (*1|Stimulus*), the condition tested (*1|Condition*), and the participants’ anonymised identifier (*1|Participant*).

In addition, we conducted simulations to compare the ability of the above model to predict preferences when the average slope for natural images varies between 0 and -4 (step: 0.05), rather than being fixed at -2.39. If there is a peak in preference at this natural slope value, we predict the best fit (lowest AIC) for models with slope values around -2.39.

**In task 3,** we tested the hypothesis of a preference for background-matching stimuli. A linear mixed-effect binomial model was run on the data. The response variable *bino_click* was defined as a binary variable where 1 indicates that the left target was chosen (i.e. clicked on) and 0 that the right target was chosen. Several explanatory variables were included, all of them exploring condition differences between the targets.

The first explanatory variable describes the level of background-matching between a target and the background. It is calculated as the difference between the two sides of the Manhattan distance between the target slope and the background slope (the background is the same for the two sides): *[abs(slope_targetL-slope_background) - abs(slope_targetR-slope_background)]*. This variable is predicted to yield a negative estimate, translating as a preference for smaller differences between a target and the background (i.e.; higher level of background-matching): if the left target slope value is closer to the background slope value than the right target, the difference will be negative while bino_click should tend towards 1 (click on left target).

To control for a general preference for natural statistics, we added the same explanatory variable as in Task 2 (*[abs(slope_targetL-2.39) - abs(slope_targetR-2.39)]*). We also included the background’s Fourier slope (*factor(background_slope);* categorical variable with three levels: -1, -2, -3), and the interaction between these two variables in case this preference is modulated by the background. As for task 2, we expect a negative estimate for the variable *[abs(slope_targetL-2.39) - abs(slope_targetR-2.39)]*, demonstrating a preference for natural scene statistics.

Finally, a last explanatory variable was the sign of the difference between the target and the background (*sign_of_target_background_difference*). It is a three-level categorical variable: ‘oo’ where the difference sign is the same for both targets, ‘+-’ where the difference between the left target and the background is positive while it is negative for the right background, and ‘-+’ where the reverse is true. A difference in the sign of the difference between the target and its background should not lead to a difference in preferences.

Three random effects were included: the stimulus displayed (*1|Stimulus*), the condition tested (*1|Condition*), and the participants’ anonymised identifier (*1|Participant*).

All continuous variables for both tasks were centred and scaled to improve the fit of the linear mixed-effect models.

All statistical analyses were performed using the software R (R Core Team, 2021, version 4.4.1). We fitted the data with linear mixed-effects models (*lme4* package, Bates et al., 2015) with REML set to false.

**Table S2:**
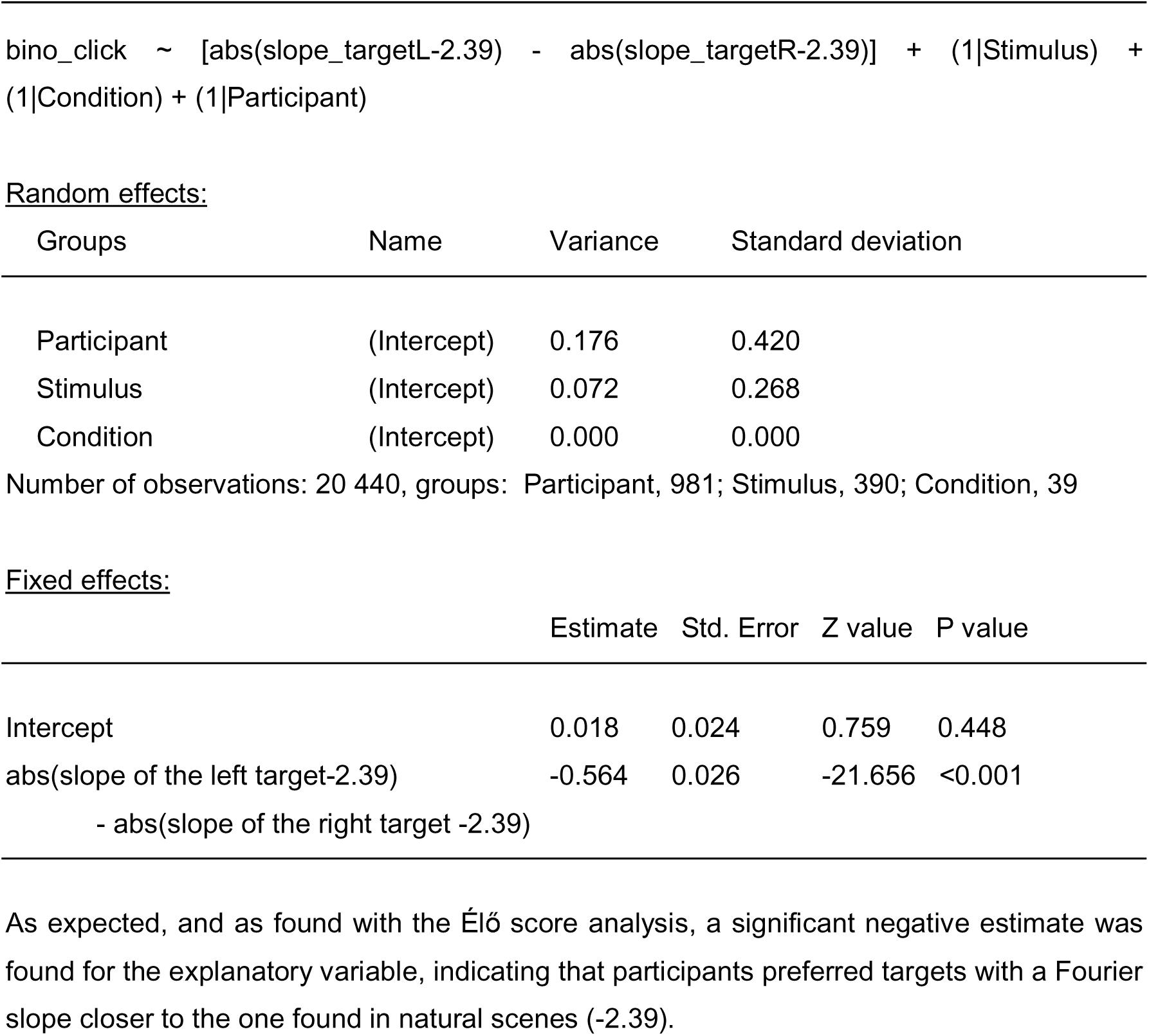
Statistical results of the binomial model for Task 2.

**Figure S3:**
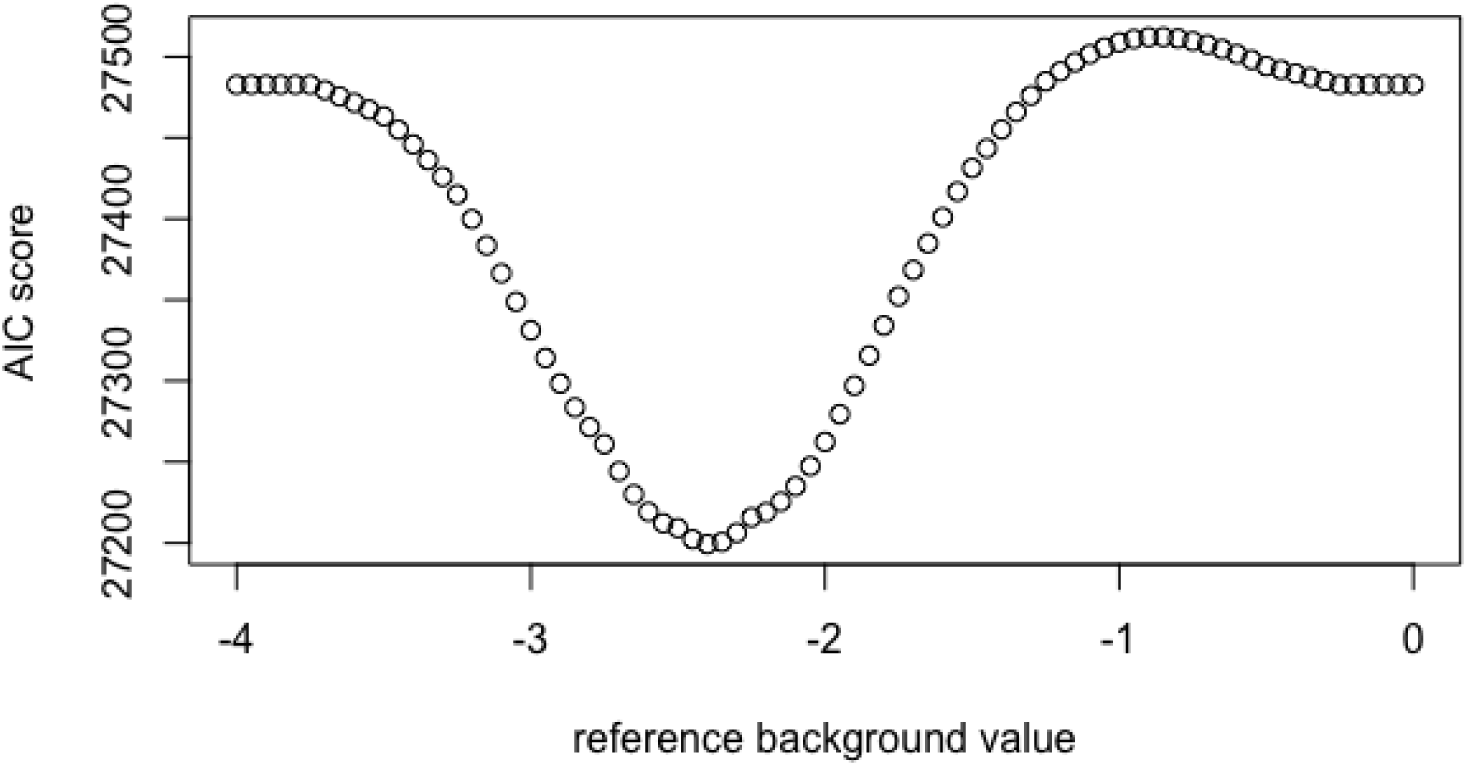
Simulation result to compare the ability of the binomial model for task 2 to predict preferences. As predicted, the lowest AIC score was found for a reference value of -2.4.

**Table S3:**
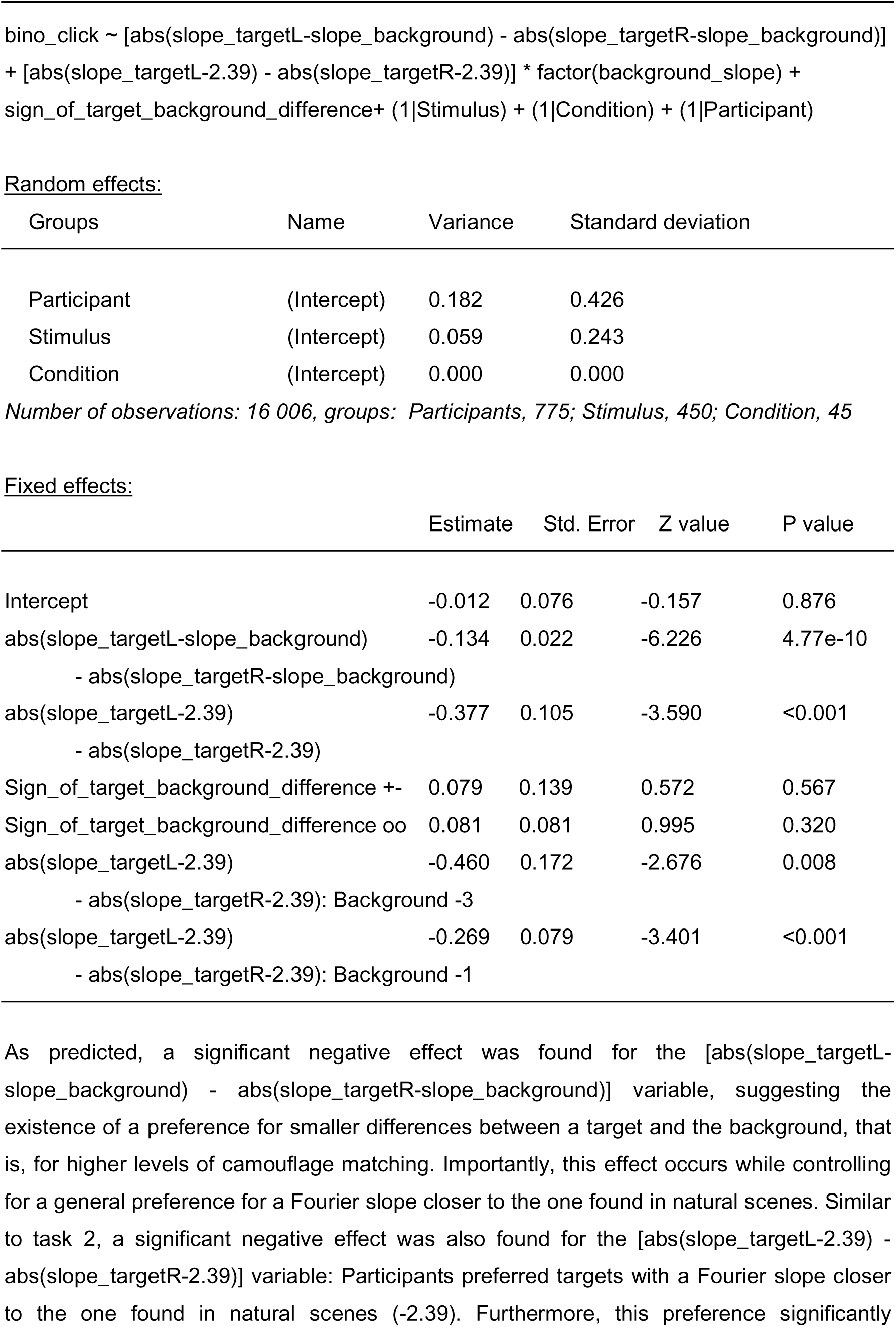
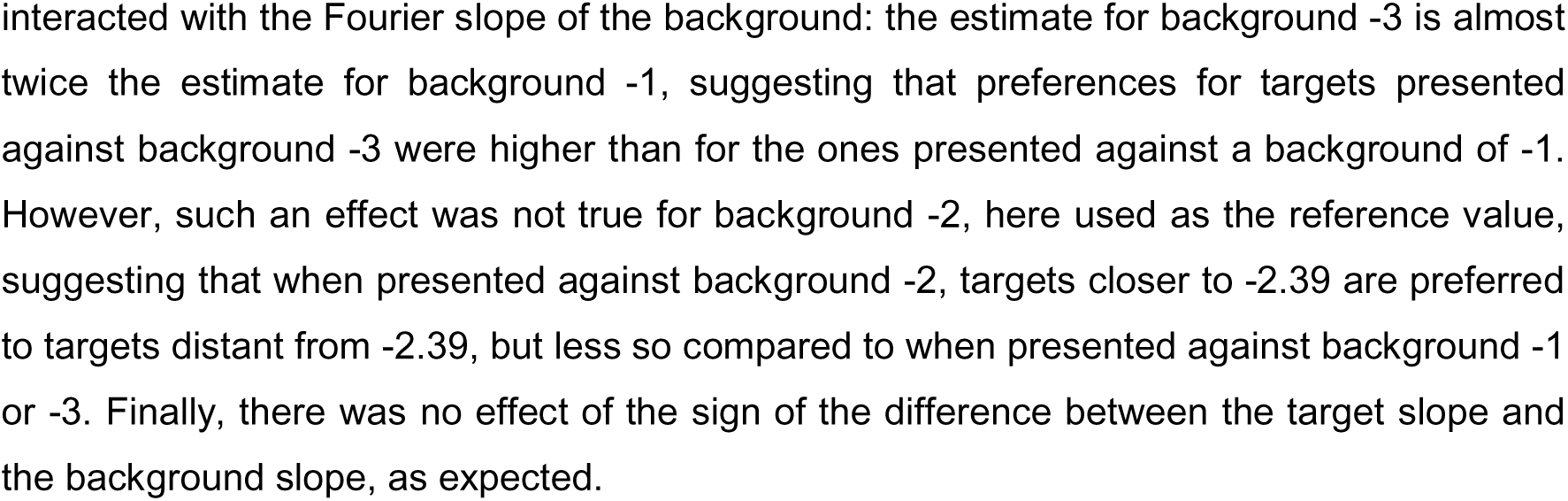
Statistical results of the binomial model for Task 3.

**Table S4:**
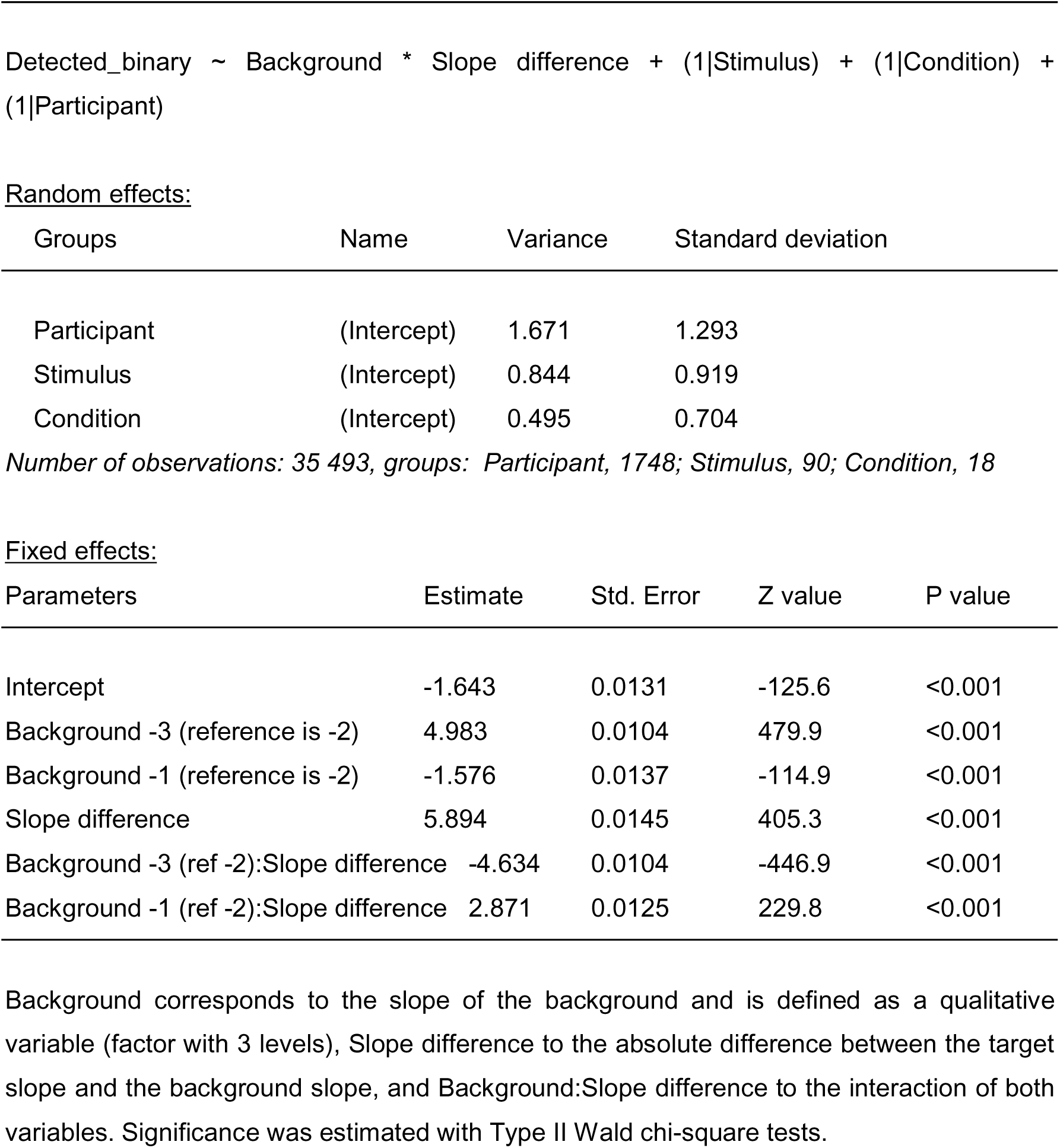
Output of a linear mixed-effects binomial model predicting detection success for Task 1.

**Figure S4.**
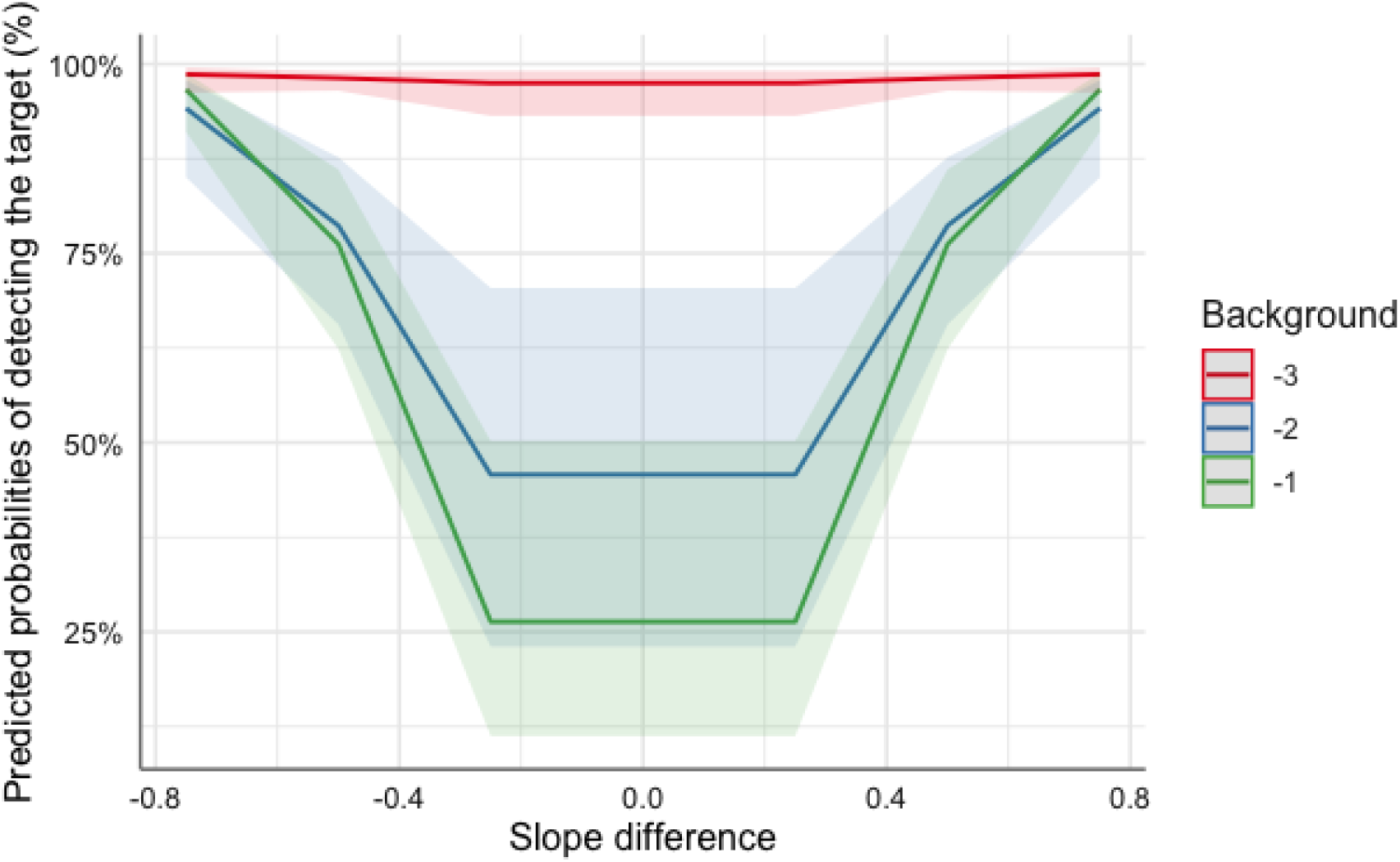
Predicted values (marginal effects) for slope difference and background values, derived from the above-mentioned binomial model (Table S4) and estimated with 95% CIs.

**Table S5:**
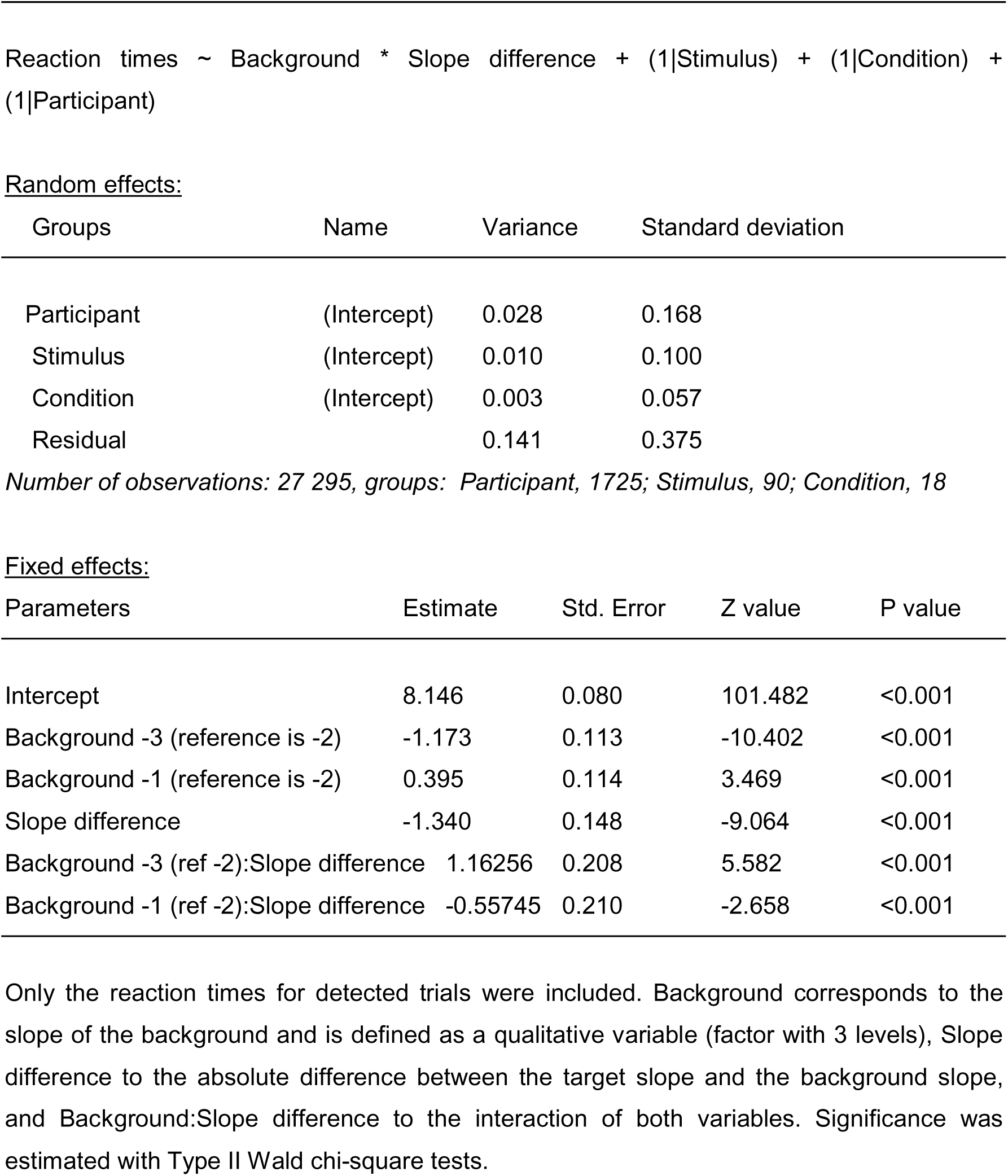
Output of the linear mixed-effects model with gamma-distributed reaction times for Task 1.

**Figure S5.**
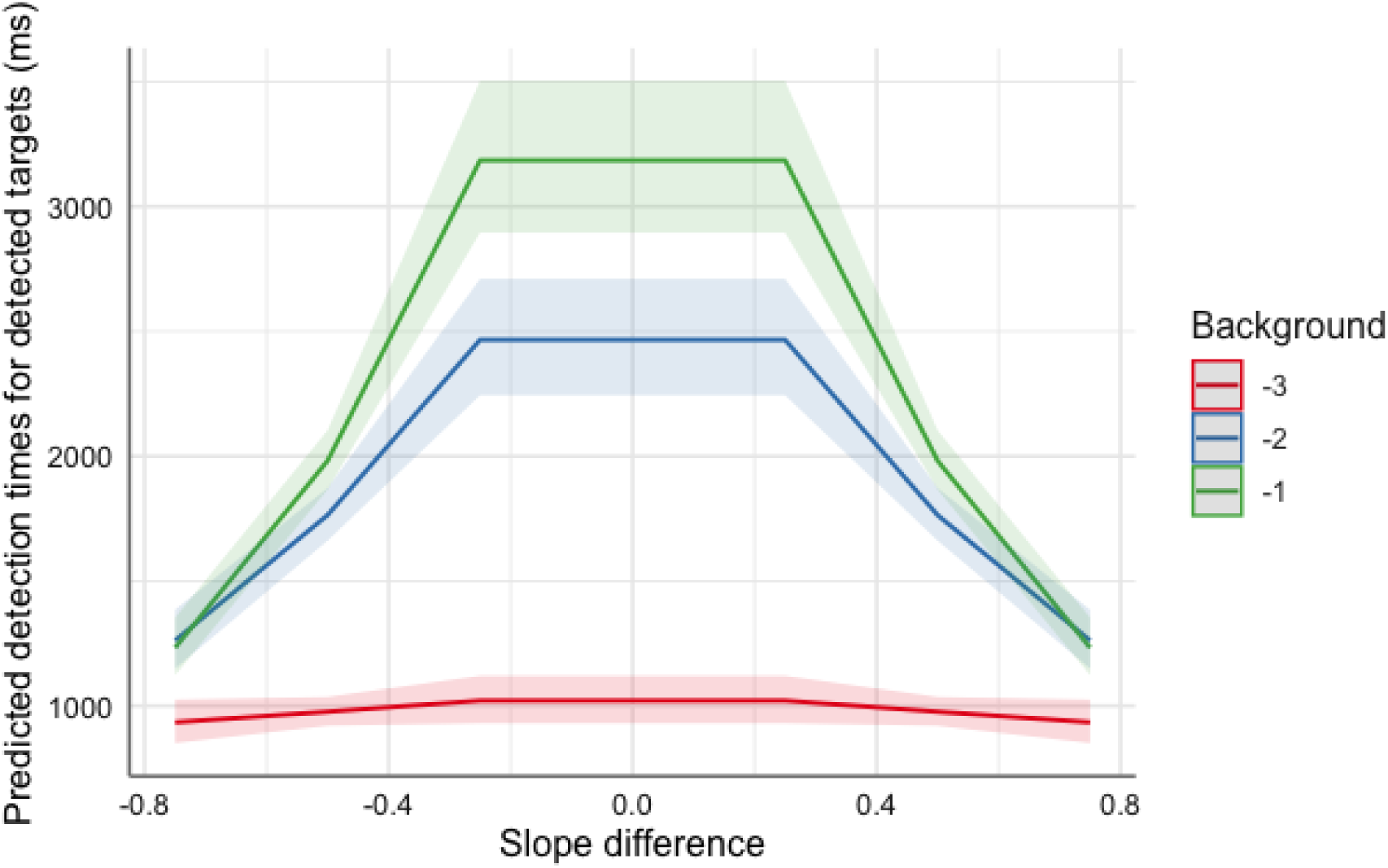
Predicted values (marginal effects) for slope difference and background values, derived from the above-mentioned model (Table S5) and estimated with 95% CIs.

**Table S6:**
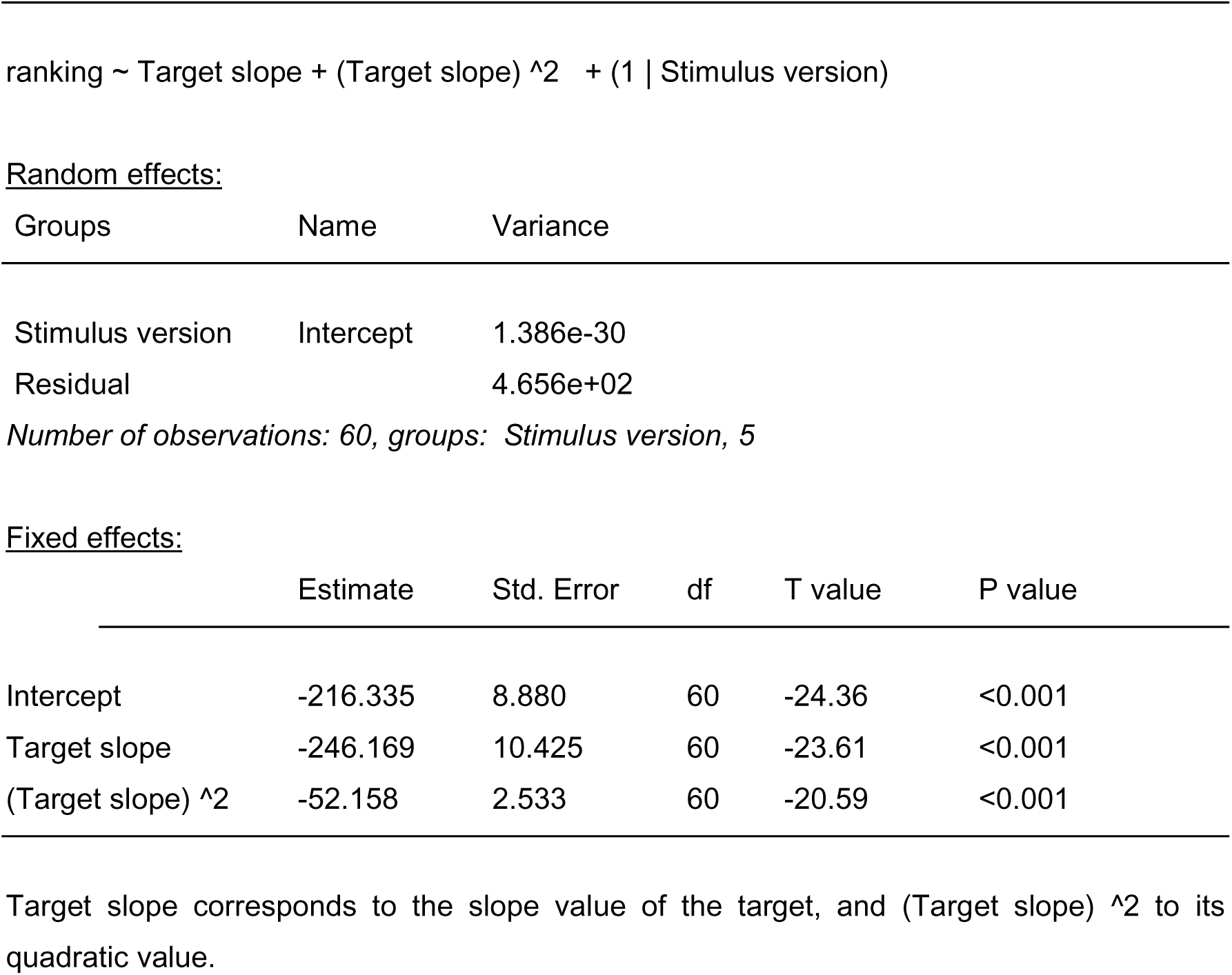
Linear mixed-effect model for predicting preferences (Élő scores) for stimuli presented against a grey background (Task 2).

**Table S7:**
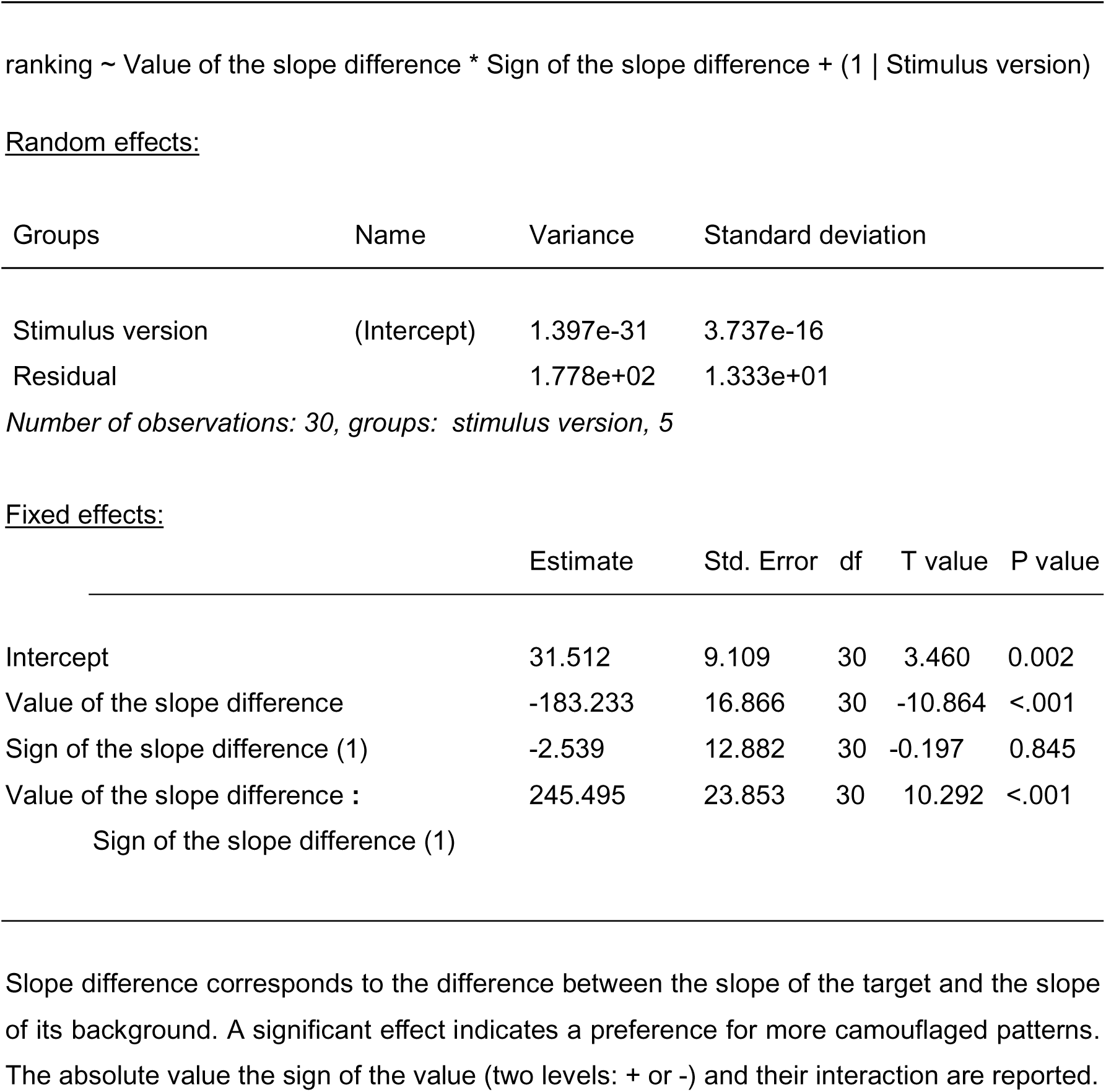
Linear mixed-effect model for predicting preferences (Élő scores) for stimuli presented against a patterned background with a Fourier slope of -1 (Task 3).

**Table S8:**
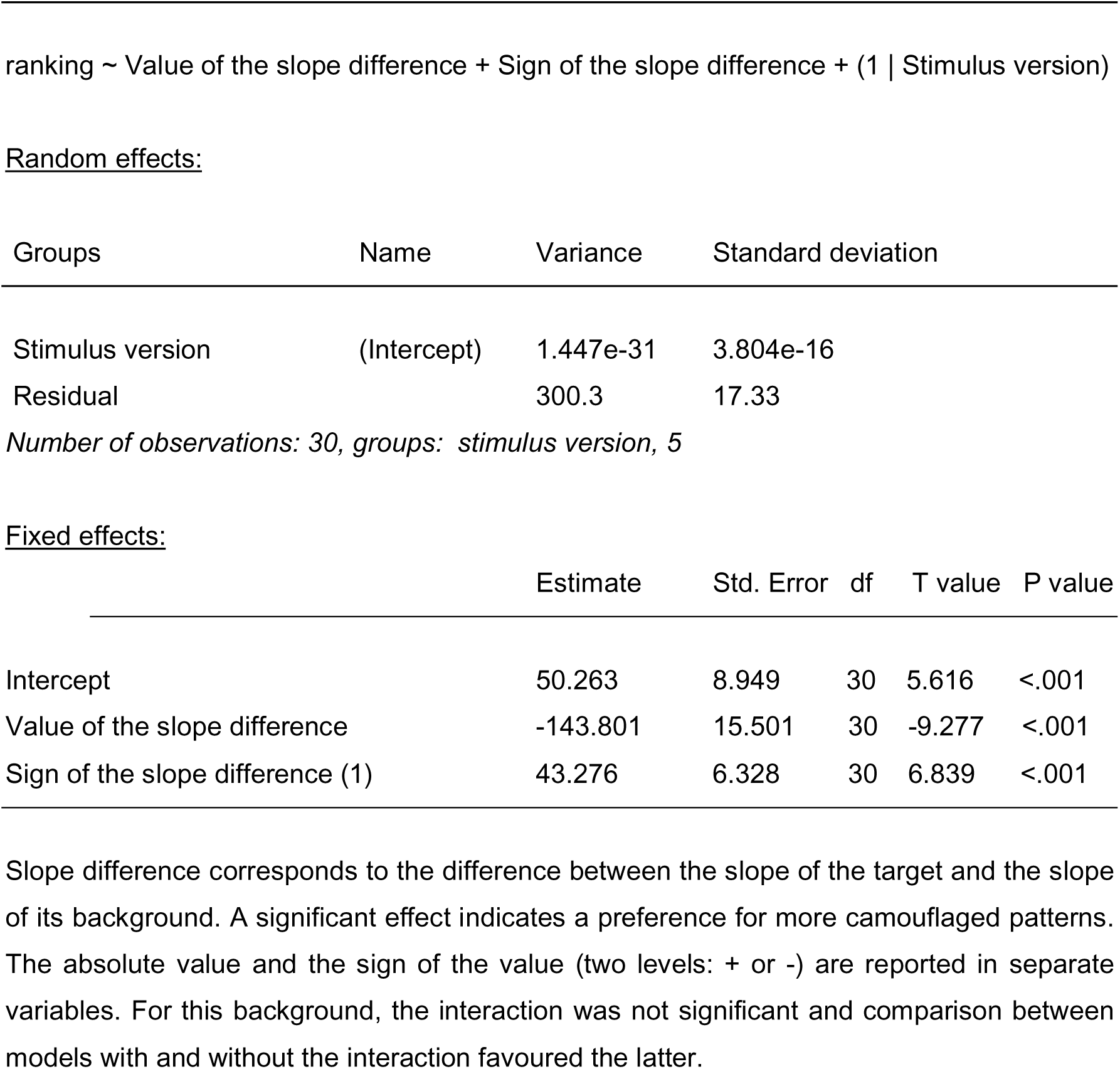
Linear mixed-effect model for predicting preferences (Élő scores) for stimuli presented against a patterned background with a Fourier slope of -2 (Task 3).

**Table S9:**
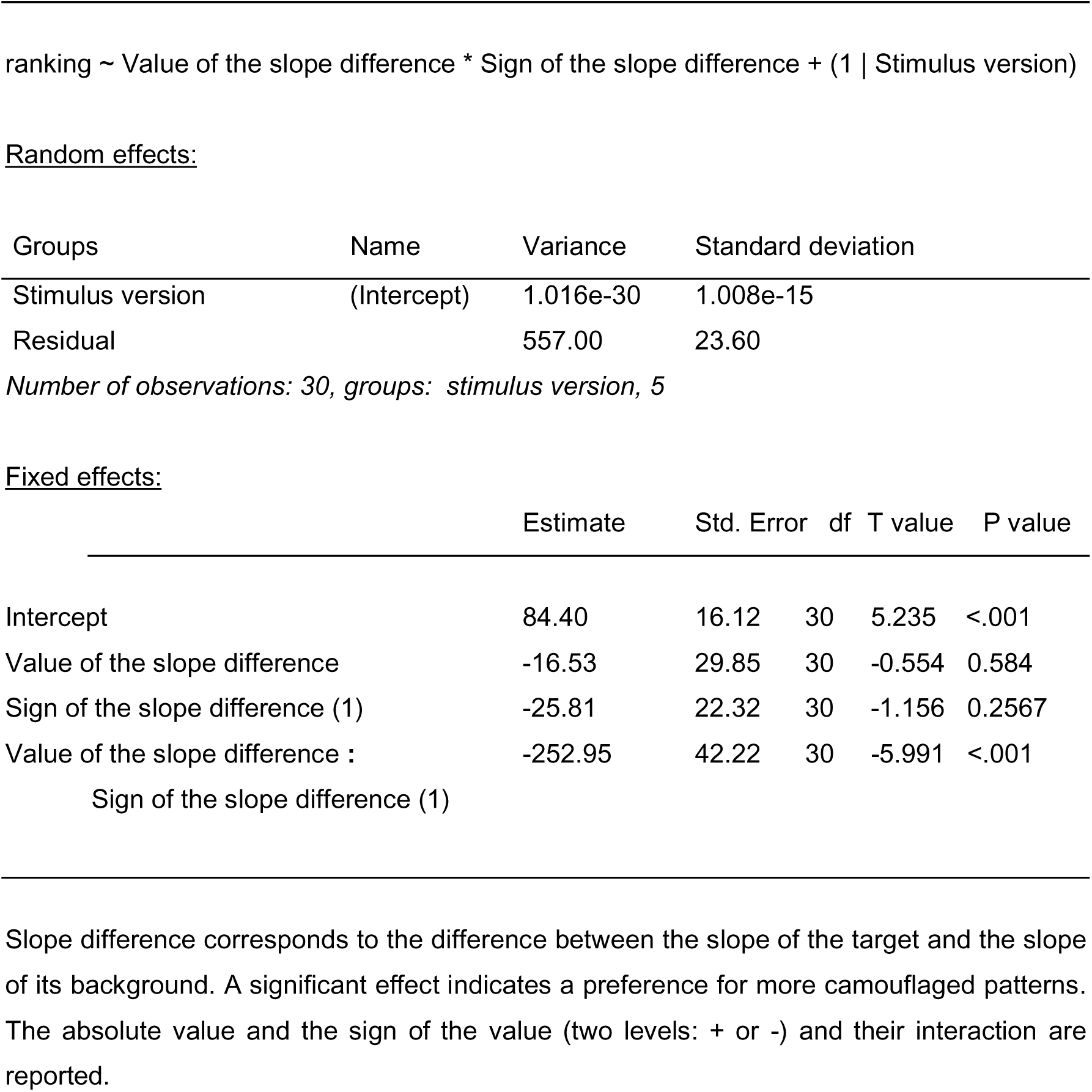
Linear mixed-effect model for predicting preferences (Élő scores) for stimuli presented against a patterned background with a Fourier slope of -3 (Task 3).

**Table S10:**
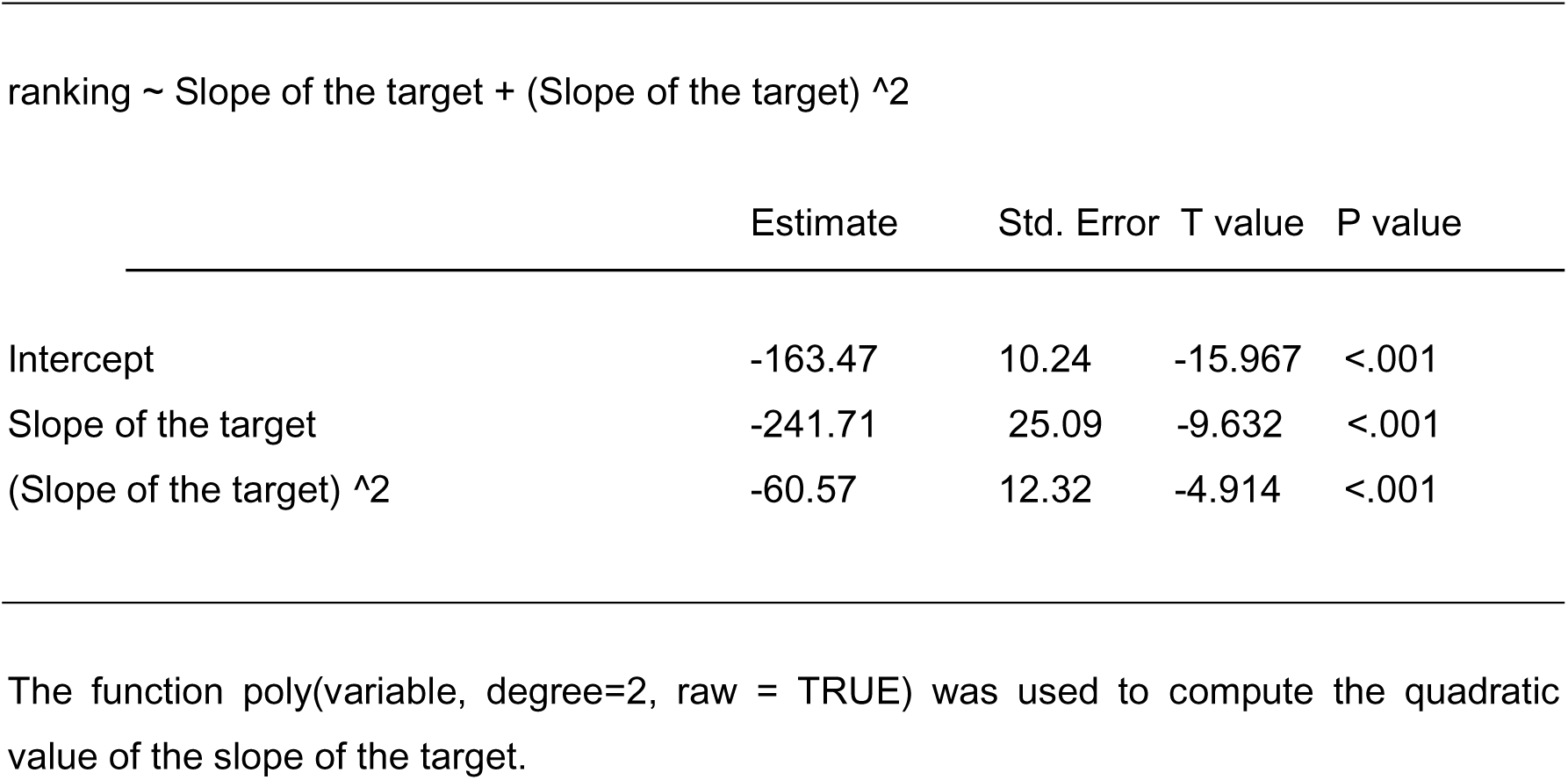
Linear quadratic model for predicting preferences (Élő scores) for stimuli presented against a patterned background with a Fourier slope of -1 (Task 3).

**Table S11:**
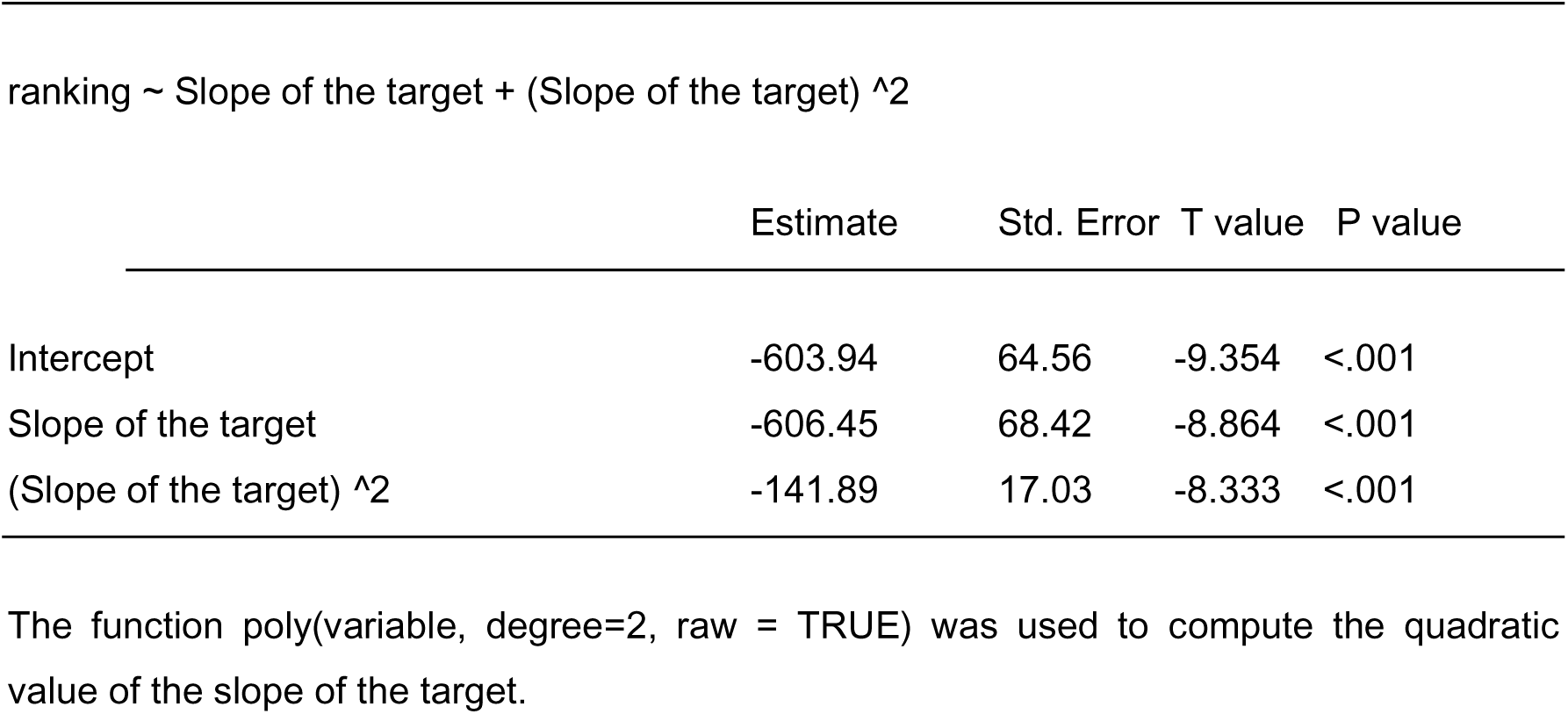
Linear quadratic model for predicting preferences (Élő scores) for stimuli presented against a patterned background with a Fourier slope of -2 (Task 3).

**Table S12:**
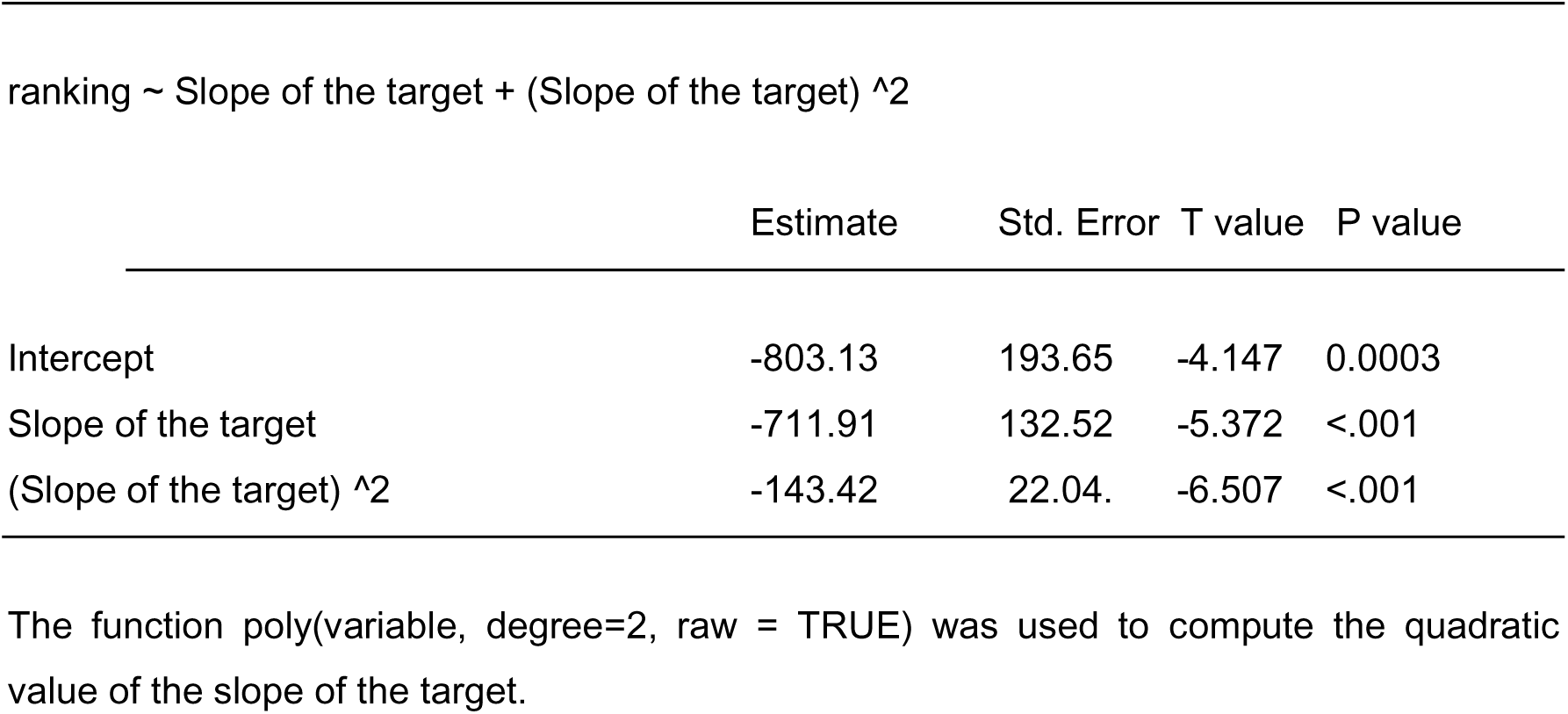
Linear quadratic model for predicting preferences (Élő scores) for stimuli presented against a patterned background with a Fourier slope of -3 (Task 3).

