## Supplementary Information for "Background-matching patterns are attractive: support for a processing bias"

¹ University of Maryland, Baltimore County, Baltimore, MD; ^2^ CEFE, University of Montpellier, CNRS, EPHE, IRD, Montpellier, France; ^3^ Department of Interdisciplinary Life Sciences, KLIVV, University of Veterinary Medicine, Vienna, Austria; ^4^ Department of Behavioural and Cognitive Biology, University of Vienna, Vienna, Austria; ^5^ Institut des Sciences de l'Evolution (CNRS-UMR 5554), Montpellier, France; ^6^ School of Biological Sciences, University of Bristol, Bristol, United Kingdom

¶ These authors contributed equally to this work

ORCID numbers:

YHB: 0000-0003-3939-3852

MR: 0000-0002-1714-6984

ICC: 0000-0002-5007-8856

TCM: 0000-0003-2938-3829

JPR: 0000-0001-6690-0085

### Supplementary material

**List of Supplementary Materials:**

Figure S1: Ratings stability for the 2-AFC task with a grey background

Figure S2: Ratings stability for the 2-AFC task with a patterned background

Table S1: Experiment questionnaire and response options

Supplementary Text S1: Participants' recruitment and procedure

Supplementary Text S2: Sociodemographic data

Supplementary Text S3: Statistical analyses with binomial models

Table S2: Statistical results of the binomial model for Task 2

Figure S3: Simulation result for Task 2

Table S3: Statistical results of the binomial model for Task 3

Table S4: Output of a linear mixed-effects binomial model predicting detection success for Task 1

Figure S4: Graphical representations of predicted fixed effects for the model presented in Table S4

Table S5: Output of the linear mixed-effects model with gamma-distributed reaction times for Task 1

Figure S5: Graphical representations of predicted fixed effects for the model presented in Table S5

Table S6: Linear mixed-effect model for predicting preferences (Élő scores) for stimuli presented against a grey background (Task 2)

Table S7: Linear mixed-effect model for predicting preferences (Élő scores) for stimuli presented against a patterned background with a Fourier slope of -1 (Task 3)

Table S8: Linear mixed-effect model for predicting preferences (Élő scores) for stimuli presented against a patterned background with a Fourier slope of -2 (Task 3)

Table S9: Linear mixed-effect model for predicting preferences (Élő scores) for stimuli presented against a patterned background with a Fourier slope of -3 (Task 3)

Table S10: Linear quadratic model for predicting preferences (Élő scores) for stimuli presented against a patterned background with a Fourier slope of -1 (Task 3)

Table S11: Linear quadratic model for predicting preferences (Élő scores) for stimuli presented against a patterned background with a Fourier slope of -2 (Task 3)

Table S12: Linear quadratic model for predicting preferences (Élő scores) for stimuli presented against a patterned background with a Fourier slope of -3 (Task 3)

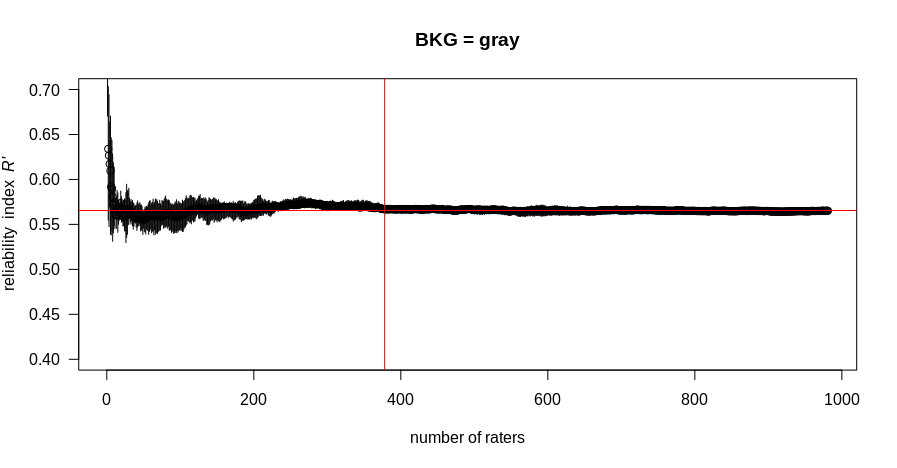

***Figure S1: Ratings stability for the 2-AFC task with a grey background.*** *The intersection of the two red lines indicates the number of raters needed to achieve the averaged reliability index (avgRI). AvgRI is measured as the average RI between 400 and 981 participants, for which we can see that ratings have achieved stability. The obtained avgRI is 0.567, corresponding to 378 raters and to a good reliability (Neumann and Clark, 2019). This was averaged across all 10 simulations, which correspond to a randomised order of raters.*

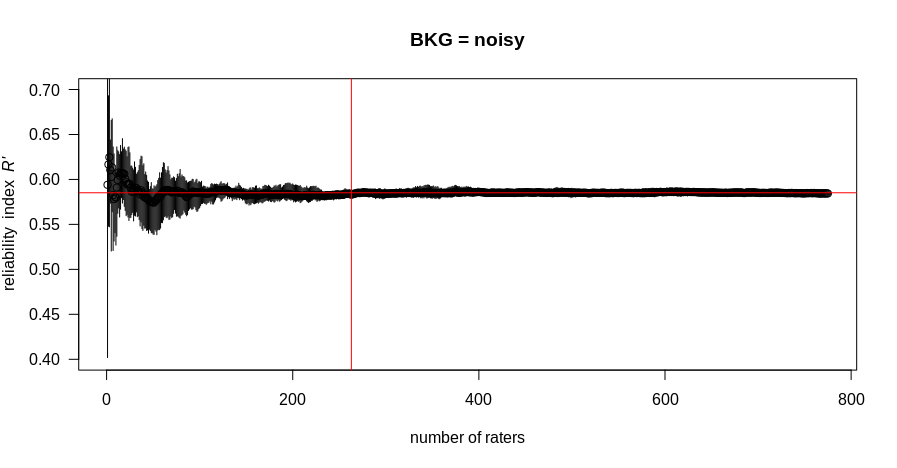

***Figure S2: Ratings stability for the 2-AFC task with a patterned background****. The intersection of the two red lines indicates the number of raters needed to achieve the averaged reliability index (avgRI). AvgRI is measured as the average RI between 250 and 775 participants, for which we can see ratings have achieved stability. The obtained avgRI is 0.585, corresponding to 263 raters and to a good reliability (Neumann and Clark, 2019). This was averaged across all 10 simulations, which correspond to a randomised order of raters.*

**Table S1: Experiment questionnaire and response options**

| *Questions* | *Options* |
| --- | --- |
| Please indicate what best describes your gender  Please indicate your birth date  Do you have uncorrected colour-blindness?  Education: select the option that corresponds to your highest qualification  What is your country of current residency?  Does your job or any of your hobbies involve visual artistic activity or visual creation?  How often do you go to art exhibitions (museums, galleries)? | Female (cis/trans), Male (cis/trans), Nonbinary, I prefer not to say, Other / I prefer to self-describe  Year and Month  Yes, No  Graduate Degree, Associate Degree or Bachelor's Degree, High School, Vocational training school certificate, CGE SSCE, PSC no diploma  List of countries  Not at all, Sometimes, Often, Daily  Once a week, Once a month, Once a year, Never |

### **Supplementary Text S1: Participants' recruitment and procedure**

**Recruitment**: Participants were recruited through advertisements on dedicated mailing lists and social networks. Participants were not paid, and they had to be at least 16 years old, corresponding to the legal digital majority in the EU. Before the experiment, participants were asked to report their month and year of birth to enforce that rule. For American citizens aged between 16 and 18 years, participants were asked to obtain their parents’ verbal approval before they participated in the experiment. Participants were asked to take part in the study only once. On top of informing them of that condition before the experiment. The IP address of the participant was only used to enforce that rule and was not associated with personal information, following a protocol approved by the CNIL. Collected data were stored on-site.

bino_click ~ [abs(slope_targetL-2.39) - abs(slope_targetR-2.39)] + (1|Stimulus) + (1|Condition) + (1|Participant)

Random effects:

Groups	Name	Variance	Standard deviation

Participant	(Intercept)	0.176	0.420

Stimulus	(Intercept)	0.072	0.268

Condition	(Intercept)	0.000	0.000

Number of observations: 20 440, groups: Participant, 981; Stimulus, 390; Condition, 39

Fixed effects:

Estimate	Std. Error	Z value	P value

Intercept	0.018	0.024	0.759	0.448

abs(slope of the left target-2.39)	-0.564	0.026	-21.656	<0.001

- abs(slope of the right target -2.39)

As expected, and as found with the Élő score analysis, a significant negative estimate was found for the explanatory variable, indicating that participants preferred targets with a Fourier slope closer to the one found in natural scenes (-2.39).

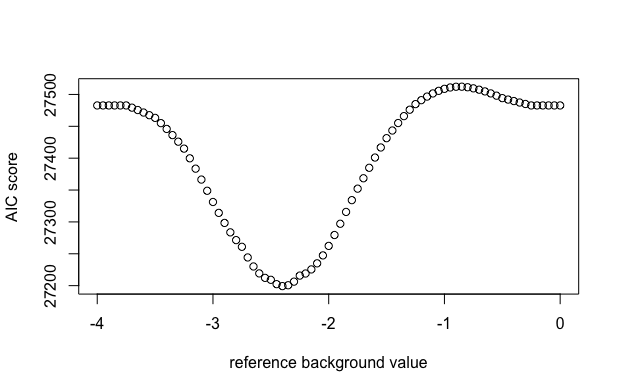

**Figure S3:** Simulation result to compare the ability of the binomial model for task 2 to predict preferences. As predicted, the lowest AIC score was found for a reference value of -2.4.

**Table S3: Statistical results of the binomial model for Task 3.**

bino_click ~ [abs(slope_targetL-slope_background) - abs(slope_targetR-slope_background)] + [abs(slope_targetL-2.39) - abs(slope_targetR-2.39)] * factor(background_slope) + sign_of_target_background_difference+ (1|Stimulus) + (1|Condition) + (1|Participant)

Random effects:

Groups	Name	Variance	Standard deviation

Participant	(Intercept)	0.182	0.426

Stimulus	(Intercept)	0.059	0.243

Condition	(Intercept)	0.000	0.000

*Number of observations: 16 006, groups: Participants, 775; Stimulus, 450; Condition, 45*

Fixed effects:

Estimate	Std. Error	Z value	P value

Intercept	-0.012	0.076	-0.157	0.876

abs(slope_targetL-slope_background)	-0.134	0.022	-6.226	4.77e-10

- abs(slope_targetR-slope_background)

abs(slope_targetL-2.39)	-0.377	0.105	-3.590	<0.001

- abs(slope_targetR-2.39)

Sign_of_target_background_difference +-	0.079	0.139	0.572	0.567

Sign_of_target_background_difference oo	0.081	0.081	0.995	0.320

abs(slope_targetL-2.39)	-0.460	0.172	-2.676	0.008

- abs(slope_targetR-2.39): Background -3

abs(slope_targetL-2.39)	-0.269	0.079	-3.401	<0.001

- abs(slope_targetR-2.39): Background -1

As predicted, a significant negative effect was found for the [abs(slope_targetL-slope_background) - abs(slope_targetR-slope_background)] variable, suggesting the existence of a preference for smaller differences between a target and the background, that is, for higher levels of camouflage matching. Importantly, this effect occurs while controlling for a general preference for a Fourier slope closer to the one found in natural scenes. Similar to task 2, a significant negative effect was also found for the [abs(slope_targetL-2.39) - abs(slope_targetR-2.39)] variable: Participants preferred targets with a Fourier slope closer to the one found in natural scenes (-2.39). Furthermore, this preference significantly interacted with the Fourier slope of the background: the estimate for background -3 is almost twice the estimate for background -1, suggesting that preferences for targets presented against background -3 were higher than for the ones presented against a background of -1. However, such an effect was not true for background -2, here used as the reference value, suggesting that when presented against background -2, targets closer to -2.39 are preferred to targets distant from -2.39, but less so compared to when presented against background -1 or -3. Finally, there was no effect of the sign of the difference between the target slope and the background slope, as expected.

**Table S4: Output of a linear mixed-effects binomial model predicting detection success for Task 1.**

Detected_binary ~ Background * Slope difference + (1|Stimulus) + (1|Condition) + (1|Participant)

Random effects:

Groups	Name	Variance	Standard deviation

Participant	(Intercept)	1.671	1.293

Stimulus	(Intercept)	0.844	0.919

Condition	(Intercept)	0.495	0.704

*Number of observations: 35 493, groups: Participant, 1748; Stimulus, 90; Condition, 18*

Fixed effects:

Parameters	Estimate	Std. Error	Z value	P value

Intercept	-1.643	0.0131	-125.6	<0.001

Background -3 (reference is -2)	4.983	0.0104	479.9	<0.001

Background -1 (reference is -2)	-1.576	0.0137	-114.9	<0.001

Slope difference	5.894	0.0145	405.3	<0.001

Background -3 (ref -2):Slope difference	-4.634	0.0104	-446.9	<0.001

Background -1 (ref -2):Slope difference	2.871	0.0125	229.8	<0.001

Background corresponds to the slope of the background and is defined as a qualitative variable (factor with 3 levels), Slope difference to the absolute difference between the target slope and the background slope, and Background:Slope difference to the interaction of both variables. Significance was estimated with Type II Wald chi-square tests.

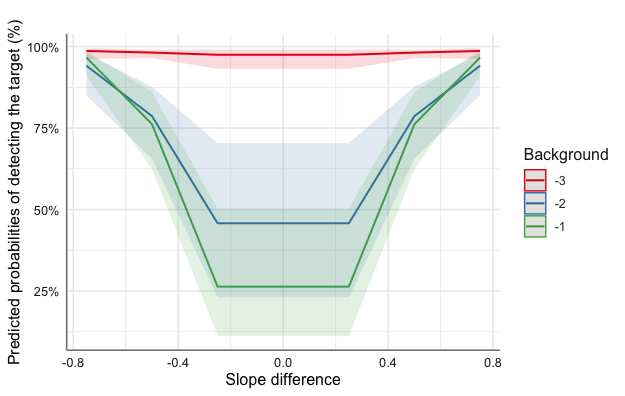

**Figure S4. Predicted values (marginal effects) for slope difference and background values**, derived from the above-mentioned binomial model (Table S4) and estimated with 95% CIs.

**Table S5: Output of the linear mixed-effects model with gamma-distributed reaction times for Task 1.**

Reaction times ~ Background * Slope difference + (1|Stimulus) + (1|Condition) + (1|Participant)

Random effects:

Groups	Name	Variance	Standard deviation

Participant	(Intercept)	0.028	0.168

Stimulus	(Intercept)	0.010	0.100

Condition	(Intercept)	0.003	0.057

Residual	0.141	0.375

*Number of observations: 27 295, groups: Participant, 1725; Stimulus, 90; Condition, 18*

Fixed effects:

Parameters	Estimate	Std. Error	Z value	P value

Intercept	8.146	0.080	101.482	<0.001

Background -3 (reference is -2)	-1.173	0.113	-10.402	<0.001

Background -1 (reference is -2)	0.395	0.114	3.469	<0.001

Slope difference	-1.340	0.148	-9.064	<0.001

Background -3 (ref -2):Slope difference	1.16256	0.208	5.582	<0.001

Background -1 (ref -2):Slope difference	-0.55745	0.210	-2.658	<0.001

Only the reaction times for detected trials were included. Background corresponds to the slope of the background and is defined as a qualitative variable (factor with 3 levels), Slope difference to the absolute difference between the target slope and the background slope, and Background:Slope difference to the interaction of both variables. Significance was estimated with Type II Wald chi-square tests.

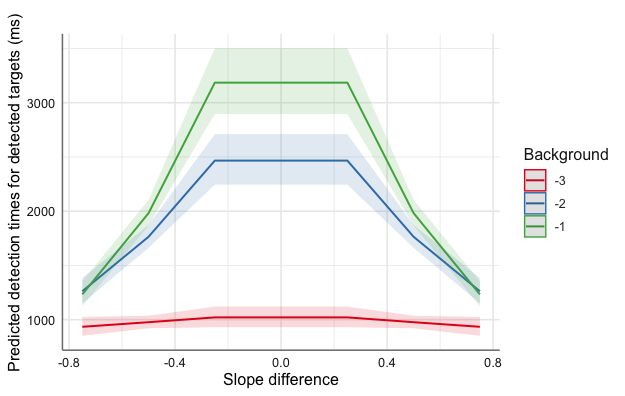

**Figure S5. Predicted values (marginal effects) for slope difference and background values**, derived from the above-mentioned model (Table S5) and estimated with 95% CIs.

**Table S6: Linear mixed-effect model for predicting preferences (Élő scores) for stimuli presented against a grey background (Task 2).**

ranking ~ Target slope + (Target slope) ^2	+ (1 | Stimulus version)

Random effects:

Groups	Name	Variance

Stimulus version	Intercept	1.386e-30

Residual	4.656e+02

*Number of observations: 60, groups: Stimulus version, 5*

Fixed effects:

Estimate	Std. Error	df	T value	P value

Intercept	-216.335	8.880	60	-24.36	<0.001

Target slope	-246.169	10.425	60	-23.61	<0.001

(Target slope) ^2	-52.158	2.533	60	-20.59	<0.001

Target slope corresponds to the slope value of the target, and (Target slope) ^2 to its quadratic value.

**Table S7: Linear mixed-effect model for predicting preferences (Élő scores) for stimuli presented against a patterned background with a Fourier slope of -1 (Task 3).**

ranking ~ Value of the slope difference * Sign of the slope difference + (1 | Stimulus version)

Random effects:

Groups	Name	Variance	Standard deviation

Stimulus version	(Intercept)	1.397e-31	3.737e-16

Residual	1.778e+02	1.333e+01

*Number of observations: 30, groups: stimulus version, 5*

Fixed effects:

Estimate	Std. Error	df	T value	P value

Intercept	31.512	9.109	30	3.460	0.002

Value of the slope difference	-183.233	16.866	30	-10.864	<.001

Sign of the slope difference (1)	-2.539	12.882	30	-0.197	0.845

Value of the slope difference **:**	245.495	23.853	30	10.292	<.001

Sign of the slope difference (1)

Slope difference corresponds to the difference between the slope of the target and the slope of its background. A significant effect indicates a preference for more camouflaged patterns. The absolute value the sign of the value (two levels: + or -) and their interaction are reported.

**Table S8: Linear mixed-effect model for predicting preferences (Élő scores) for stimuli presented against a patterned background with a Fourier slope of -2 (Task 3).**

ranking ~ Value of the slope difference + Sign of the slope difference + (1 | Stimulus version)

Random effects:

Groups	Name	Variance	Standard deviation

Stimulus version	(Intercept)	1.447e-31	3.804e-16

Residual	300.3	17.33

*Number of observations: 30, groups: stimulus version, 5*

Fixed effects:

Estimate	Std. Error	df	T value	P value

Intercept	50.263	8.949	30	5.616	<.001

Value of the slope difference	-143.801	15.501	30	-9.277	<.001

Sign of the slope difference (1)	43.276	6.328	30	6.839	<.001

Slope difference corresponds to the difference between the slope of the target and the slope of its background. A significant effect indicates a preference for more camouflaged patterns. The absolute value and the sign of the value (two levels: + or -) are reported in separate variables. For this background, the interaction was not significant and comparison between models with and without the interaction favoured the latter.

**Table S9: Linear mixed-effect model for predicting preferences (Élő scores) for stimuli presented against a patterned background with a Fourier slope of -3 (Task 3).**

ranking ~ Value of the slope difference * Sign of the slope difference + (1 | Stimulus version)

Random effects:

Groups	Name	Variance	Standard deviation

Stimulus version	(Intercept)	1.016e-30	1.008e-15

Residual	557.00	23.60

*Number of observations: 30, groups: stimulus version, 5*

Fixed effects:

Estimate	Std. Error	df	T value	P value

Intercept	84.40	16.12	30	5.235	<.001

Value of the slope difference	-16.53	29.85	30	-0.554	0.584

Sign of the slope difference (1)	-25.81	22.32	30	-1.156	0.2567

Value of the slope difference **:**	-252.95	42.22	30	-5.991	<.001

Sign of the slope difference (1)

Slope difference corresponds to the difference between the slope of the target and the slope of its background. A significant effect indicates a preference for more camouflaged patterns. The absolute value and the sign of the value (two levels: + or -) and their interaction are reported.

**Table S10: Linear quadratic model for predicting preferences (Élő scores) for stimuli presented against a patterned background with a Fourier slope of -1 (Task 3).**

ranking ~ Slope of the target + (Slope of the target) ^2

Estimate	Std. Error T value	P value

Intercept	-163.47	10.24	-15.967	<.001

Slope of the target	-241.71	25.09	-9.632	<.001

(Slope of the target) ^2	-60.57	12.32	-4.914	<.001

The function poly(variable, degree=2, raw = TRUE) was used to compute the quadratic value of the slope of the target.

**Table S11: Linear quadratic model for predicting preferences (Élő scores) for stimuli presented against a patterned background with a Fourier slope of -2 (Task 3).**

ranking ~ Slope of the target + (Slope of the target) ^2

Estimate	Std. Error T value	P value

Intercept	-603.94	64.56	-9.354	<.001

Slope of the target	-606.45	68.42	-8.864	<.001

(Slope of the target) ^2	-141.89	17.03	-8.333	<.001

The function poly(variable, degree=2, raw = TRUE) was used to compute the quadratic value of the slope of the target.

**Table S12: Linear quadratic model for predicting preferences (Élő scores) for stimuli presented against a patterned background with a Fourier slope of -3 (Task 3).**

ranking ~ Slope of the target + (Slope of the target) ^2

Estimate	Std. Error T value	P value

Intercept	-803.13	193.65	-4.147	0.0003

Slope of the target	-711.91	132.52	-5.372	<.001

(Slope of the target) ^2	-143.42	22.04. -6.507	<.001

The function poly(variable, degree=2, raw = TRUE) was used to compute the quadratic value of the slope of the target.
